# Targeting endothelial FOXO1 protects diabetic β-cells and improves wound healing

**DOI:** 10.1101/2023.12.04.569948

**Authors:** Hao Wu, Kui Cui, Bo Zhu, Yao Wei Lu, Liang Sun, Qianyi Ma, Ashish Jian, Daniel Osorio Hurtado, Xinlei Gao, Yanqian Li, Sudarshan Bhattacharjee, Haojie Fu, Amy E. Birsner, Marina V. Malovichko, Manna Li, Beibei Wang, Bandana Singh, Qianman Peng, Scott Wong, Vikram Norton, Kathryn Li, Shahram Eisa-Beygi, Donghai Wang, Douglas B. Cowan, Diane R. Bielenberg, Sanjay Srivastava, Shira Rockowitz, Jian Xu, Robert J. D’Amato, Kaifu Chen, Min Dong, Sudha Biddinger, Hong Chen

## Abstract

The forkhead box O1 (FOXO1) transcription factor plays critical roles in regulating not only metabolic activity but also angiogenesis in the vascular endothelium^1–4^. Our previous studies show that epsin endocytic adaptors can regulate both angiogenesis and lymphangiogenesis^5–7^. Endothelial cells (ECs) lining the inside of blood vessels are continuously exposed to circulating insulin and insulin-like growth factors (IGFs). Emerging evidences suggest that ECs can affect β-cell function^8–11^. Excessive IGF2, especially elevated local IGF2 levels in islets, may represent a risk factor for developing diabetes^12–15^; however, the underlying molecular mechanisms by which aberrant angiogenesis and endothelium-derived factors regulate pancreatic β-cell function in diabetes remain unclear. Here, we report that the pancreas of diabetic patients as well as the pancreas, skin, and plasma of streptozotocin/high fat diet (STZ/HFD)-induced diabetic mice and *db/db* mice contains excess IGF2, which can lead to β-cell dysfunction and apoptosis. Single-cell transcriptomics combined with mass spectrometry analysis reveal that endothelial-specific knockout of FOXO1 increases circulating soluble and cell-membrane or intracellular expression levels of IGF type 2 receptor (IGF2R) and CCCTC-binding factor (CTCF), while decreasing IGF2 levels in diabetes. Both IGFR2^15–17^ and CTCF^18–21^ can reduce IGF2 levels and may ameliorate β-cell decline associated with excess IGF2 in diabetes. Furthermore, depletion of FOXO1, epsins, or knockdown of ULK1 inhibits autophagy formation in ECs, preventing degradation of vascular endothelial growth factor receptor 2 (VEGFR2) to promote angiogenesis and improve wound healing in diabetes. Our findings reveal that endothelial FOXO1 regulates epsin-dependent angiogenesis and affects β-cell function and fate through CTCF and IGF2-IGF2R, providing a potential strategy for ameliorating diabetes and accelerating cutaneous wound healing.

## Introduction

Inadequate supply of functional insulin due to the loss of pancreatic β-cells results in diabetes^22^. The regenerative potential of pancreatic islet β-cells is extremely restricted; therefore, the massive loss of β-cells in diabetes necessitates the urgent development of new therapeutic strategies^23^. Pancreatic endothelial cells play a crucial role in supporting pancreatic β-cells by providing oxygen to metabolically active cells, alongside mediating glucose input and insulin output^8,11^. Moreover, the cells of the islet vasculature regulate β-cell activity through the secretion of growth factors and other molecules^8,10^. Despite these critical roles, the precise mechanisms by which endothelial cells (ECs) directly influence β-cell function and viability in diabetes remain unclear.

The transcription factor forkhead box O1 (FOXO1) plays a critical role in regulating vascular growth by coordinating metabolic and proliferative activities in ECs^1,3^. The EC-specific deletion of FOXO1 in mice results in a marked increase in EC proliferation, while overexpression of FOXO1 limits vascular expansion, resulting in vessel thinning and hypobranching^1^. Epsins, as a family of evolutionarily conserved adaptor proteins, play critical roles in endocytosis^5^. Our previous studies have revealed the interaction between epsins and VEGFR2/3 regulates angiogenesis and lymphangiogenesis, wound healing, as well as the pathogenesis of diabetes^6,7,24^. Sespite this, the molecular mechanisms underlying the regulation of epsin-dependent angiogenesis in diabetes by FOXO1 signaling remain unclear.

Insulin-like growth factor 2 (IGF2) was detected in diabetic foot ulcers and surrounding tissues^25^. A novel genetic variant in the IGF2 gene is associated with a 20% reduced risk for type 2 diabetes (T2D)^14^. Transgenic mice overexpressing IGF2 specifically in β-cells can develop T2D^26^. Functionally, IGF2 renders islets more susceptible to β-cell damage and autoimmune attack^13^. At least two studies have delineated that methylation of a CCCTC-binding factor (CTCF)-dependent boundary governs the expression of IGF2^18,19^. Furthermore, IGF type 2 receptor (IGF2R) serves as a scavenger for circulating IGF2^15–17,27^. However, in the context of diabetes, whether FOXO1 cooperates with CTCF in endothelial cells to regulate β-cell function and viability through the IGF2-IGF2R system has yet to be determined.

Cell-identity switches have been identified in both humans and animals^28–30^. Influence of age on β-cells was reconstituted from heterologous islet cells after near-total β-cell loss in diabetic mice upon the combined action of FOXO1 and downstream effectors^31^. The single-cell RNA sequencing (scRNA-seq) analysis of pseudo islets revealed that human α- and γ-cells have the potential to be reprogrammed into glucose-dependent insulin secretors, under diabetic conditions^32^. Single-cell transcriptomics of the human endocrine pancreas revealed that IGF2 expression levels are significantly increased in β-cells of T2D patients^33^. Pseudotime analysis of stem-cell-derived β-cells (SC-β) indicates that *Igf2* expression decreases to maintain β-cell identity^34^. Despite recent advancements, our understanding of pancreatic islet cell plasticity remains limited due to a lack of a blueprint of single-cell pancreas transcriptomic data under different diabetic conditions.

To explore the impact of pancreatic ECs on β-cell function and survival and to elucidate the underlying molecular regulatory mechanisms, we generated a comprehensive pancreatic single-cell atlas using pancreas from mice, including wild-type (WT) mice, as well as WT and EC-specific knockout FOXO1 or epsin mice administered streptozotocin (STZ) and high-fat diet (HFD). Accordingly, we sought to gain insight into the cell-cell communications between ECs and β-cells in addition to the molecular mechanisms that contribute to improved β-cell function and enhanced wound healing in diabetes.

## Results

### FOXO1 upregulates epsins and ULK1 in ECs

Chromatin immunoprecipitation with sequencing (ChIP-seq) analysis show that FOXO1 can directly bind to the promoter regions of *EPN1* and *ULK1* respectively in human umbilical vein endothelial cells (HUVECs), indicating FOXO1 can upregulate the expression of both epsins 1 and 2 as well as ULK1^3^. The acetylation of H3K27 and the trimethylation of H3K4 indicated an active state for FOXO1 bound to *EPN1* and *ULK1* (**Fig. 1a, b**). Furthermore, luciferase and ChIP assays revealed that FOXO1 is a *bona fide* target of *Epn1* and *Ulk1* in mouse primary pancreatic ECs (**Extended Fig. 1a-d**).

**Fig. 1.**
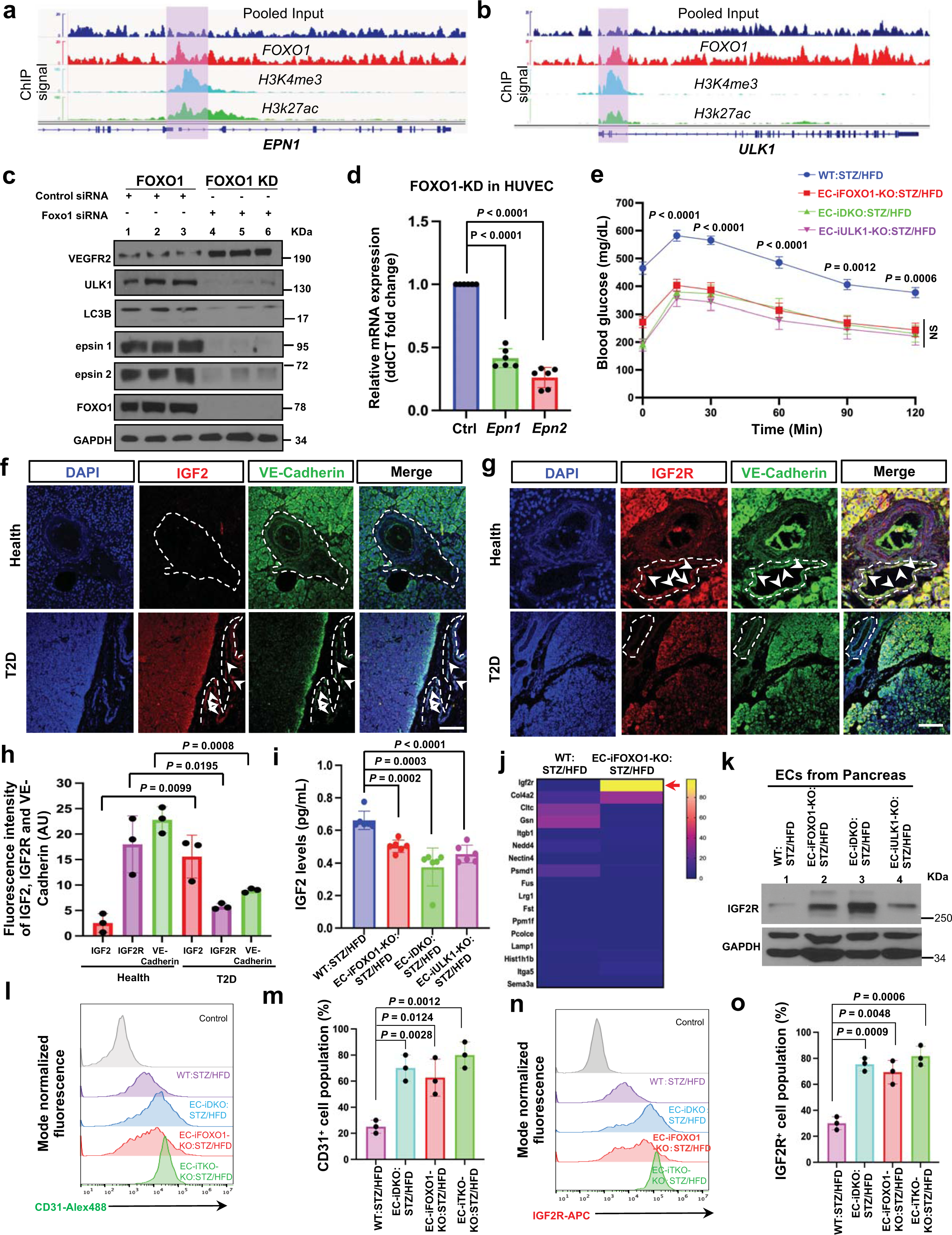
EC-specific knockout of FOXO1, epsins, and ULK1 ameliorates diabetes, increases IGF2R expression while reducing IGF2 levels in diabetes. **a, b**, ChIP-Seq signals in HUVECs for FOXO1, the histone modifications H3k4me3, H3k27ac at the genomic loci of (**a**) *EPN1,* (**b**) *ULK1*. FOXO1 exhibits high binding signal peaks at the *EPN1* and *ULK1* promoter regions, while H3K4me3 and H3K27ac also show elevated peaks at these sites (highlighted with pink boxes), suggesting that FOXO1 may upregulate epsin 1 and ULK1 in EC. Track range FOXO1, input, 0-20; H3K4me3, H3K27ac, 0-100. ChIP-seq signals are expressed in reads per kilobase million (RPKM). **c**, Western blot analysis for the protein levels of FOXO1, epsins 1 and 2, ULK1, LC3B, ULK1, VEGFR2 in the normal and *FOXO1* knockdown HUVECs (n = 3). **d**, qRT-PCR analysis of *EPN1* and *EPN2* expression in control and *FOXO1* knockdown HUVECs (n = 6). **e**, GTT analysis showed that EC-specific knockout of FOXO1, epsins, and ULK1 in diabetic mice resulted in decreased fasting blood glucose levels after administrated 2.5 g/kg 10% glucose compared with WT:STZ/HFD mice (n = 10). **f, g**, Immunofluorescence detection of IGF2 (red, f) and IGF2R (red, g) along with VE-Cadherin (green, **f, g**) in the pancreas of healthy individuals (Top) and T2D patients (bottom). **h**, Quantification of IGF2, IGF2R, and VE-Cadherin signal intensities in (**f**) and (**g**). Representative results from n = 3 healthy and T2D patients, respectively. **i**, ELISA of IGF2 in blood plasma from WT:STZ/HFD, EC-iFOXO1-KO:STZ/HFD, EC-iDKO:STZ/HFD, EC-iULK1:STZ/HFD mice (n = 6 per group). **j**, Heatmap of mass spectrometry data illustrating factors in the supernatants of ECs isolated from WT:STZ/HFD and EC-iFOXO1-KO:STZ/HFD mice, incubated in the EC-conditioned medium (n = 3). **k**, Western blot analysis for IGF2R in the ECs of pancreas from the WT:STZ/HFD, EC-iFOXO-KO:STZ/HFD, EC-iDKO:STZ/HFD, and EC-iULK1-KO:STZ/HFD mice (n = 3). **l-o**, Flow cytometry analysis of CD31^+^ cells (**l**) and quantification (**m**); IGF2R^+^ cells (**n**) and quantification (**o**) in the pancreas from WT:STZ/HFD, EC-iFOXO1-KO:STZ/HFD, EC-iDKO:STZ/HFD, and EC-iTKO:STZ/HFD (EC-triple knockout of epsins and ULK1) mice (n = 3 per group). Statistical tests in **d, i, m, o**, significance was assessed by one-way ANOVA (*P* < 0.0001 overall) with post hoc Dunnett’s test, compared to WT:STZ/HFD control mice; **e**, two-way ANOVA (*P* < 0.0001 overall) followed by Bonferroni’s multiple comparison test, comparisons for EC-iFOXO1-KO:STZ/HFD vs WT:STZ/HFD; **h**, two-way ANOVA followed by Bonferroni’s multiple comparison test. Data in represent mean ± S.E.M. NS, not significant. Scale bars, 50 μm (**f, h**).

Of note, *FOXO1* knockdown (**Extended data Fig. 1e**) shows significantly decreased mRNA levels of *Epns 1* and *2* (**Fig. 1d**), *ULK1, LC3B, BECN1* (**Extended data Fig. 1f**) and increased *KDR* levels (**Extended data Fig. 1g**) compared to normal in HUVECs. Subsequently, FOXO1 knockdown increased VEGFR2 expression but decreased protein levels of epsins 1 and 2, ULK1, and LC3B (**Fig. 1c**).

### EC-FOXO1 depletion improves β-cell function and ameliorates diabetes

Earlier research has demonstrated that FOXO1 regulates angiogenesis^1,3,35^ and β-cell function in diabetes^1,3,31,36^. We previously showed that epsins negatively regulate angiogenesis^5,6^ and lymphangiogenesis in diabetes^7^. Ash *et al.* reported that autophagy mediates VEGFR2 degradation, leading to impaired angiogenesis^37,38^. Given that FOXO1 upregulates epsins and ULK1(**Fig. 1a– d**, **Extended Data Fig. 1a–d, f**), we sought to explore the mechanisms by which ECs regulate β-cell function in diabetes. To achieve this, we generated endothelial-specific knockouts for either FOXO1, epsins, or ULK1, in diabetic mice. The mice were treated with streptozotocin (STZ) and maintained on a high-fat diet (HFD). All mice (hereafter referred to as WT:STZ/HFD or WSH, EC-iFOXO1-KO:STZ/HFD or FSH, EC-iDKO:STZ/HFD or DSH, EC-iULK1-KO:STZ/HFD) exhibited symptoms similar to type 2 diabetes, with no significant difference in weight gain. However, EC-iFOXO1-KO:STZ/HFD, EC-iDKO:STZ/HFD, and EC-iULK1-KO:STZ/HFD mice showed improved glucose tolerance by the glocuse tolerance tests (GTT) (**Fig. 1e**), insulin sensitivity by the insulin tolerance test (ITT) (**Extended Data Fig. 1h**), and decreased total cholesterol and triglyceride levels (**Extended Data Fig. 1i, j)** compared to WT:STZ/HFD mice, indicating that endothelial-specific knockout of either FOXO1, epsins, or ULK1 ameliorates diabetes. Glucose-stimulated insulin secretion (GSIS) assays revealed that when islets isolated from EC-iFOXO1-KO:STZ/HFD, EC-iDKO:STZ/HFD, and EC-iULK1-KO mice were treated with a high concentration of 25 mM glucose for 2 hours, β-cell insulin secretion was significantly increased compared with no glucose treatment. However, in WT:STZ/HFD mice, insulin secretion levels were significantly reduced after 25 mM glucose treatment compared with none glucose treatment. Notably, when islets isolated from these knockout mice were treated with a high concentration of 25 mM glucose, insulin secretion levels were significantly higher than in WT:STZ/HFD mice treated with the same high concentration of glucose (**Extended Data Fig. 1k-m)**. These results illustrate that EC-specific depletion of either FOXO1, epsins or ULK1 enhances β-cell function and mitigates diabetes, respectively.

### IGF2 and IGF2R expression in human

IGF2 overexpression leads to β-cell dysfunction and enhances susceptibility to damage, ultimately resulting in the development of T2D in transgenic mice^13,26^. Moreover, elevated levels of IGF2 in the circulation of individuals with diabetes and obesity are associated with β-cell apoptosis, which is further supported by a meta-analysis of loss-of-function variants in IGF2 from GWAS studies^13,14,26,39^. However, the detailed molecular mechanisms are unclear. To dissect the mechanisms linking diabetes, obesity, IGF2-IGF2R system, and β-cell apoptosis, we first examined the expression levels of IGF2 and IGF2R in pancreatic tissues obtained from individuals diagnosed with T2D as well as from healthy individuals. Notably, the pancreatic tissue obtained from individuals with T2D showed increased levels of IGF2 (**Fig. 1f, h**) and decreased expression of IGF2R (**Fig. 1g, h**) compared to healthy individuals, indicating dysregulation of the IGF2-IGF2R system in pancreatic tissue may contribute to the development of T2D.

### IGF2 and IGF2R expression in diabetic mice

Next, we investigated the levels of IGF2 in the plasma of endothelial-specific knockouts of FOXO1 diabetic mice. Conspicuously, plasma IGF2 levels exhibited a significant increase in WT:STZ/HFD mice compared to EC-iFOXO1-KO:STZ/HFD mice (**Fig. 1i**). Given that FOXO1 upregulates epsins and ULk1 (**Fig. 1a–d**, **Extended Data Fig. 1a–d, f**), we sought to determine whether plasma IGF2 levels were decreased in diabetic mice lacking epsins or ULK1 in EC. Consistently, plasma IGF2 levels in EC-iDKO:STZ/HFD and EC-iULK1-KO:STZ/HFD mice were significantly reduced compared to those in WT:STZ/HFD mice (**Fig. 1i**). These findings suggest that FOXO1, epsins, and ULK1 in EC may play a key role in regulating IGF2 levels in diabetes.

To elucidate whether FOXO1 regulates the molecular mechanism of diabetic complications through the IGF2-IGF2R system, we performed mass spectrometry (MS) assay on proteins in the serum-free medium supernatant of pancreatic ECs purified from diabetic WT:STZ/HFD, EC-iFOXO1-KO:STZ/HFD, EC-iDKO:STZ/HFD, and EC-iULK1-KO:STZ/HFD mice. Notably, in EC-iFOXO1-KO:STZ/HFD mice compared to WT:STZ/HFD mice, there was a nearly 100-fold increase in the levels of IGF2R, a scavenger for circulating IGF2^15–17,27^. This suggests that the EC-specific deletion of FOXO1 leads to a marked increase of soluble IGF2R levels under diabetic conditions (**Fig. 1j**).

Immunofluorescent analysis of the pancreas tissues showed higher levels of IGF2R and VE-cadherin, along with increased co-localization of both proteins in EC-iFOXO1-KO:STZ/HFD, EC-iDKO:STZ/HFD, and EC-iULK1-KO:STZ/HFD mice compared to WT:STZ/HFD mice (**Extended Data Fig. 2a-c**). Furthermore, western blot and immunofluorescence analyses showed that IGF2R protein levels in pancreatic ECs isolated from EC-iFOXO1-KO:STZ/HFD, EC-iDKO:STZ/HFD, and EC-iULK1-KO:STZ/HFD mice were significantly increased compared with those in ECs from pancreas of WT:STZ/HFD mice (**Fig. 1k**, **Extended Data Fig. 2d-h**). These observations were confirmed by flow cytometry analysis of whole pancreatic cells. Notably, flow cytometry of whole pancreatic cells revealed a significant increase of approximately 50% in the CD31**^+^** cell population (**Fig. 1l, m**) and about 60% increase in the IGF2R**^+^** cell population **(Fig. 1n, o)** in EC-iFOXO1-KO:STZ/HFD, EC-iDKO:STZ/HFD mice compared to WT:STZ/HFD mice. Notably, diabetic mice with EC triple knockout of epsins 1, 2 and ULK1 (EC-iTKO:STZ/HFD) exhibited a significant increase in CD31^+^ cells and IGF2R^+^ cells compared with WT:STZ/HFD (**Fig. 1l-o**). These findings indicated that endothelial-specific knockout of FOXO1, epsins, or ULK1 increases expression levels of IGF2R in both circulating soluble forms and endothelial cell membrane-associated forms.

**Fig. 2.**
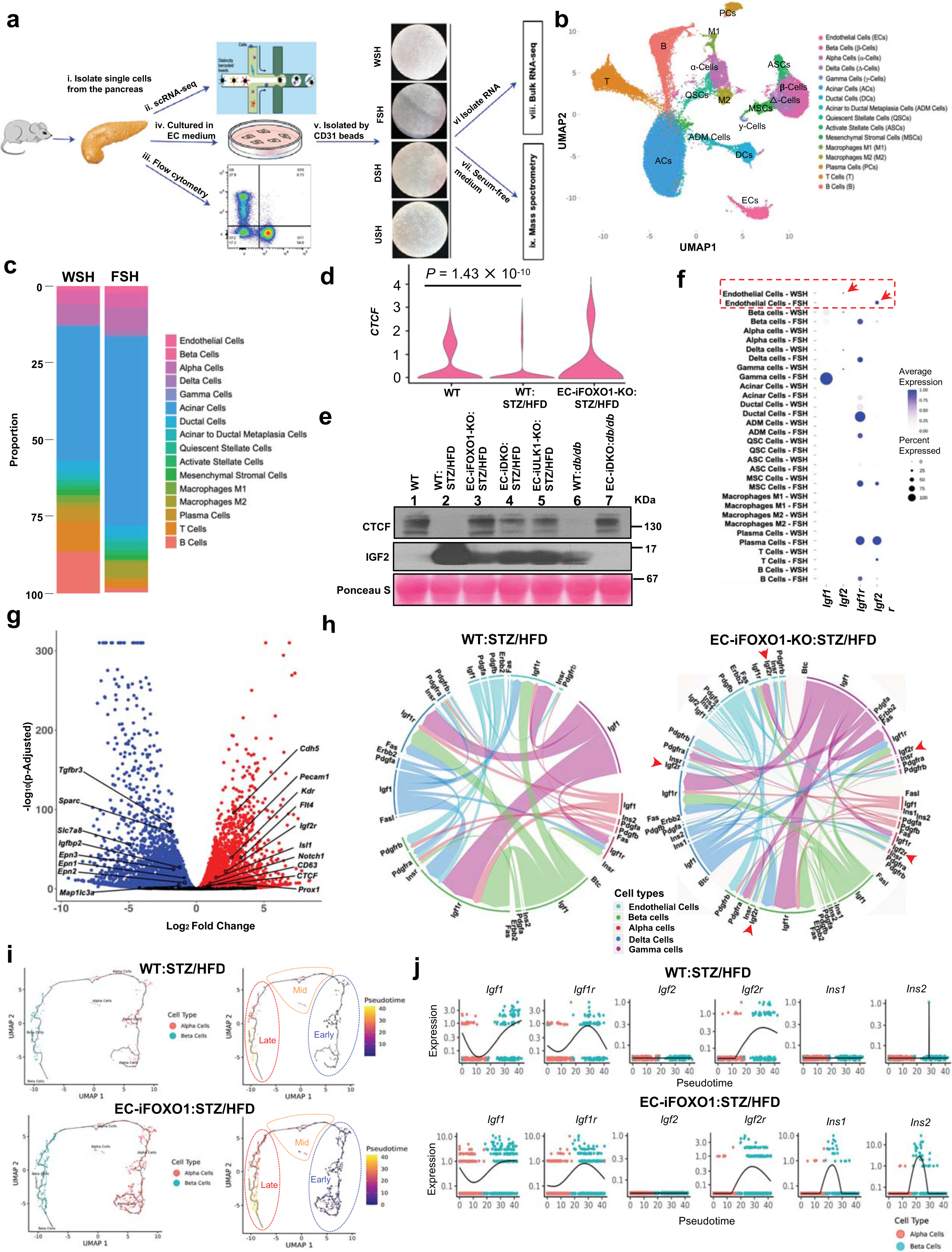
scRNA-seq and bulk RNA-seq analyses reveal that the endothelial cell-specific knockout of FOXO1 increases the expression and activities of IGF2R and CTCF, while decreasing IGF2 levels in diabetes. **a**, Schematic of the data generation and study design: Isolated single cells from mouse pancreas (i), Single cell RNA-seq (ii), Flow cytometry (iii), EC purification (iv and v), Bulk RNA-seq (viii), and Mass spectrometry (ix) of pancreas from the WT:STZ/HFD, EC-iFOXO1-KO:STZ/HFD, EC-iDKO:STZ/HFD, and EC-iULK-KO/STZ/HFD mice. **b**, The pancreas scRNA-seq UMAP plot displays 16 clusters, with cells colored according to their assigned clusters. **c**, Cell-type proportions analysis showed an increase in both ECs and β-cells in EC-iFOXO-KO:STZ/HFD (FSH) compared with WT:STZ/HFD (WSH) mice. **d, e**, Violin plot (**d**) shows that *CTCF* expression is significantly reduced in ECs from WT:STZ/HFD compared with WT mice. However, *CTCF* was significantly increased in EC-iFOXO1-KO:STZ/HFD mice compared with WT:STZ/HFD mice. Western blot (**e**) analysis of mouse plasma showed reduced CTCF levels and increased IGF2 levels in WT:STZ/HFD mice compared with WT mice. However, in the plasmas of EC-iFOXO1-KO:STZ/HFD, EC-iDKO:STZ/HFD, and EC-iULK1-KO:STZ/HFD mice, CTCF was increased while IGF2 decreased compared with WT:STZ/HFD mice. Consistently, EC-iDKO:*db/db* mice had increased CTCF and decreased IGF2 in plasma compared with WT:*db/db* mice (n = 3). **f**, Dot plots depict the expression levels of the genes *Igf1*, *Igf2*, *Igf1r*, and *Igf2r* for each cluster. *Igf2r* levels were significantly increased in ECs from EC-iFOXO1-KO:STZ/HFD mouse pancreas compared with ECs from WT:STZ/HFD mouse pancreas. However, the expression of *Igf2* was significantly reduced in EC-iFOXO1-KO:STZ/HFD pancreatic EC compared with WT:STZ/HFD, as shown by the red dotted box. **g**, Volcano plot of bulk RNA-seq showing the differentially expressed genes (DEGs) from ECs between EC-iFOXO1-KO:STZ/HFD and WT:STZ/HFD. Genes, including *Igf2r*, *CTCF*, *Kr*, and *CD63* were upregulated in EC-iFOXO1-KO:STZ/HFD compared with WT:STZ/HFD. The genes were color-coded based on fold change (>1.5-fold) and FDR-p values (p < 0.05). Statistical analysis employed the two-sided Wilcoxon test implemented via Seurat. **h**, Interaction between ECs and β-cells by using NichNet reveals links were established between prioritized ligands from ECs (senders) and target genes in islets endocrine cells, including β-cells (receivers) in pancreases from EC-iFOXO1-KO:STZ/HFD versus WT:STZ/HFD. The thickness of the ribbons is proportional to the ligand’s regulatory potential, and ribbon color indicates the subcluster of origin for each ligand. IGF2R signaling (red arrowhead) is significantly more active in EC-iFOXO-KO:STZ/HFD pancreases compared to WT:STZ/HFD. **i, j**, In silico pseudotime ordering of α-cells (red) to β-cells (blue) reveals three distinct states along the main pseudotemporal trajectory: early, mid, and late. Each dot represents an individual cell, with the majority aligning along the main path from early to late, indicative of reprogramming progression (**i**). The expression of selected genes is illustrated along the pseudotime from α-cells to β-cells. Among these genes, except for *Ins1* and *Ins2*, *IGF2R* exhibits a significantly increased expression in EC-iFOXO1-KO:STZ/HFD compared to WT:STZ/HFD along the pseudotime trajectory from α-to-β cells (**j**).

Taken together, our data indicates that in diabetic mice, the specific knockout of FOXO1, epsins, or ULK1 in ECs leads to notable alterations in the expression levels of IGF2 and IGF2R.

### EC-FOXO1 depletion increases pancreatic EC and β-cell proportions in diabetic mice

To better understand and build a detailed map of pancreatic ECs and β-cells, their interactions under diabetic conditions and the underlying cellular and molecular mechanisms, we conducted single-cell RNA sequencing (scRNA-seq) on pancreatic samples from WT, WT:STZ/HFD, EC-iFOXO1-KO:STZ/HFD, EC-iDKO:STZ/HFD, EC-iULK1-KO:STZ/HFD mice (**Fig. 2**, **Fig. 3h**, **Extended Data Fig. 3, 4a, b**). We also performed bulk RNA sequencing of ECs isolated from these mice (**Fig. 2g**, **Extended Data Fig. 5**). After filtering and quality control, scRNA-seq produced a total of 13,933 profiles with an average of 4,776 UMIs per cell. Overall, 18,461 distinct genes were detected in more than 3 cells, and 2000 highly variable genes were selected for data dimension reduction. Subsequently, 16 clusters were annotated based on the basis of known lineage marker genes and visualized as a uniform manifold approximation and projection (UMAP) plot **(Fig. 2b)**.

**Fig. 3.**
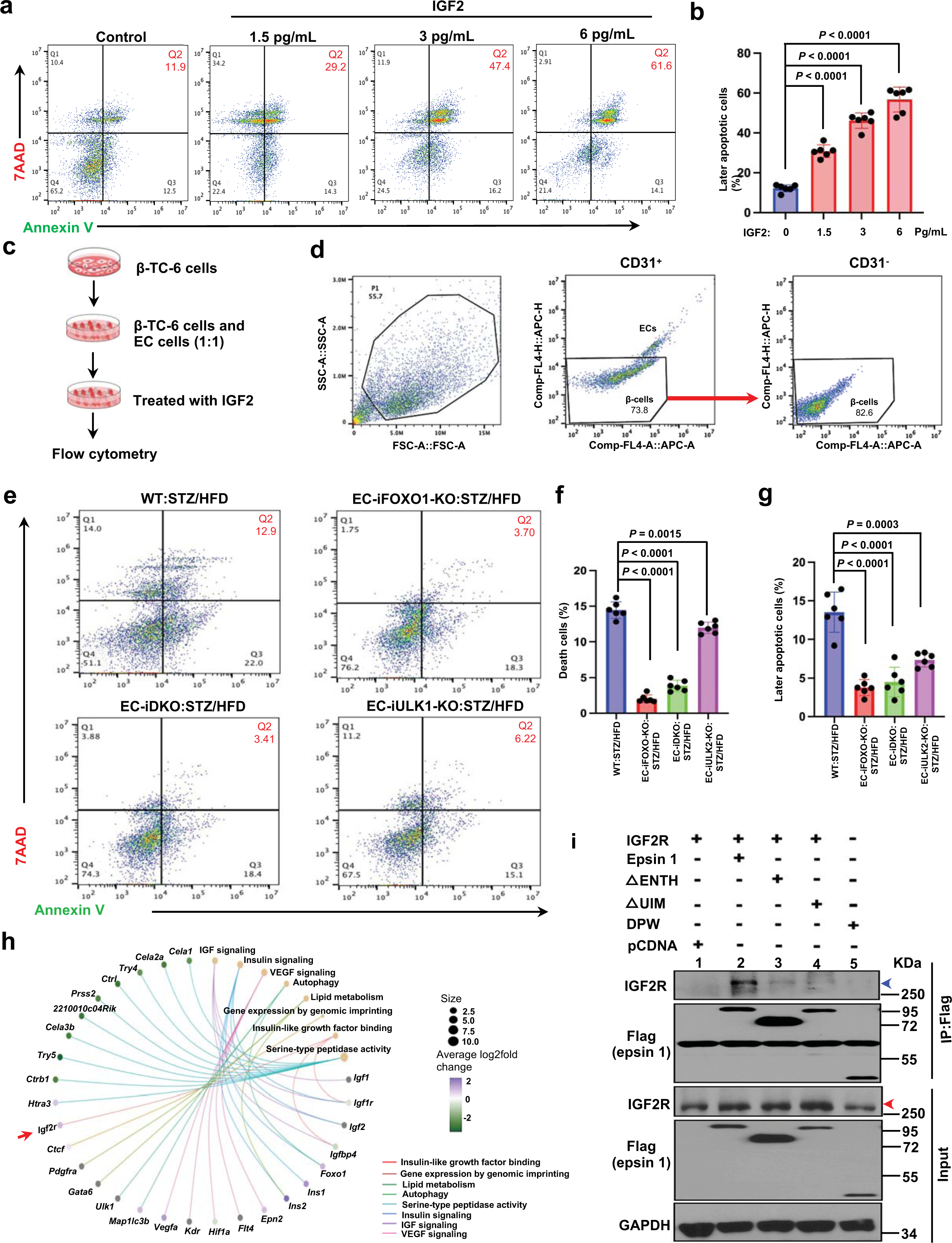
EC-specific knockout FOXO1, epsins, ULK1 prevent β-cell apoptosis through the IGF2-IGF2R system in diabetes. **a**, Flow cytometry analysis of apoptosis of β-cells (β-TC-6) induced by IGF2. Representative FACS plot and the percentage of Q2 shows Annexin V-positive and 7-AAD-positive populations, which are cells undergoing later apoptosis. **b**, Quantification of percentage of later apoptosis of β-TC-6 cells reveals a dose-dependent manner after treated with IGF2 at 1.5, 3, 6 Pg/mL for 24 h in **a** (n = 6). **c**, Schematic of apoptosis or death analysis of β-TC-6 cells cocultured with ECs and treated with IGF2. **d**, Gating strategy: CD31^-^ cells were β-TC-6 cells, CD31^+^ cells were ECs isolated from pancreases of WT:STZ/HFD, EC-iFOXO1-KO:STZ/HFD, EC-iDKO:STZ/HFD, and EC-iULK1-KO:STZ/HFD. Only the CD31^-^ were gated for following cell death and later apoptosis assay. **e**, Representative FACS plots of β-TC-6 cells (CD31^-^) and the percentages of Q1 (death cells, Annexin V-positive and 7-AAD-positive populations), and Q2 (later apoptosis). **f, g**, Quantification of percentages of later apoptosis (Q1) (**f**) and death cells (Q2) (**g**) in e. (n = 6). **h**, Gene concept network diagram showing correspondingly enriched gene ontology (GO) terms based on differentially expressed genes (DEGs) between EC-iFOXO1-KO:STZ/HFD mice and WT:STZ/HFD mice. i, Immunoprecipitation indicating full-length and domain-deleted constructs of FLAG-epsin 1 expressed in HEK 293T cells by endogenously enriched expressed IGF2R from the mouse primary EC cells of EC-iFOXO1-KO:STZ/HFD (n = 3). Statistical analyses in **b, f, g**, were one-way ANOVA (P < 0.0001 overall) with Tukey’s multiple comparisons test, compared with WT:STZ/HFD control mice. Data in represent mean ± S.E.M.

**Fig. 4.**
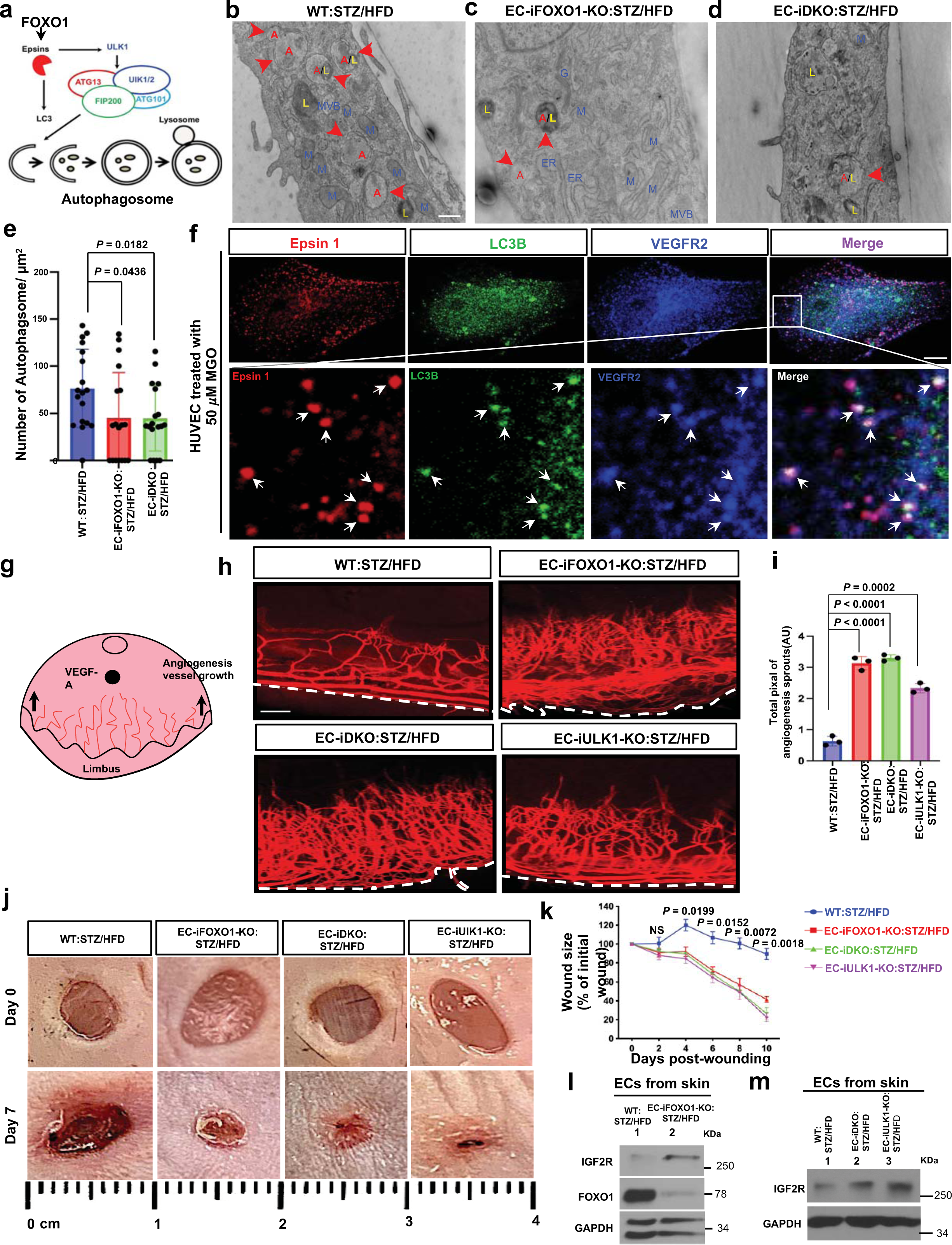
EC-specific knockout of FOXO1, epsins, and ULK1 improves wound healing by increasing angiogenesis in diabetes. **a**, Schematic showing that the FOXO1-epsins axis regulates ULK1 and LC3B to promote autophagosome formation. **b-d**, Representative electron microscopy (EM) images show electron-dense vacuoles of double membrane autophagosome (arrowheads, red) in the primary ECs of pancreases isolated from WT:STZ/HFD (**b**), EC-iFOXO1-KO:STZ/HFD (**c**), EC-iDKO:STZ/HFD (**d**) mice. L, Lysosome, M, Mitochondria, MVB, Multivesicular bodies (n = 18). **e**, Quantification of autophagosome in (**b-d**). **f,** Representative confocal immunofluorescent images of colonization epsin 1, LC3B and VEGFR2, as shown by white arrows (n = 3). **g**, Schematic showing of corneal micropocket for pellet implantation. The implanted pellets of the vessel growth factor VEGF-A can induce blood vessel growth from the limbus. **h**, Representative images of mouse corneas from WT:STZ/HFD, EC-iFOXO1-KO:STZ/HFD, EC-iDKO:STZ/HFD, and EC-iULK1-KO:STZ/HFD implanted with VEGF-A or buffer containing pellets and stained for PECAM1. White dotted line shows the limbus of mouse corneas. **i**, Quantification of PECAM1^+^ vessel area per corneas from WT:STZ/HFD, EC-iFOXO1-KO:STZ/HFD, EC-iDKO:STZ/HFD, and EC-iULK1-KO:STZ/HFD mice in (**h**) (n=5). **j**, Representative images of dermal wound healing 0, and 7 days after dermal biopsy of WT:STZ/HFD, EC-iFOXO1-KO:STZ/HFD, EC-iDKO:STZ/HFD, EC-iULK1-KO:STZ/HFD mice (n = 5). **k**, Quantification of wound area shown in (j) and presented as wound healing curve. Quantifications by using NIH Image J. **l, m**, Western blot analysis show increased IGF2R levels in skin ECs isolated from EC-iFOXO1-KO:STZ/HFD (**l**), EC-iDKO:STZ/HFD and EC-iULK1-KO:STZ/HFD (**m**) compared with skin ECs from WT:STZ/HFD (n = 3). Statistical analyses in **e, i** were one-way ANOVA with Tukey’s multiple comparisons test, compared with WT:STZ/HFD control mice. Significance in **j** was two-way ANOVA followed by Bonferroni’s multiple comparison test, comparisons for EC-iFOXO1-KO:STZ/HFD vs WT:STZ/HFD. Data in represent mean ± S.E.M. NS, not significant. Scale bars: 500nm (**b-d**), 100 μm (**h**).

A recent study identified CD63 as a potential marker of a specific subset of β-cells that exhibit high expression levels of CD63^40^. This subset of β-cells plays a pivotal role in insulin secretion and the maintenance of glucose homeostasis in both mouse models and humans with T2D^40^. Consistently, our scRNA-seq study also revealed significantly higher expression of *CD63* in β-cells of normal WT mice compared to diabetic WT:STZ/HFD mice **(Extended Data Fig. 3b, c)**. Notably, bulk RNA-seq analysis also revealed that *CD63* was upregulated in ECs from EC-iFOXO1-KO:STZ/HFD and EC-iDKO:STZ/HFD mice when compared to WT:STZ/HFD mice **(Fig. 2g**, **Extended data Fig. 5a, g)**. Our findings were further supported by a study conducted by Wang *et al.*, which revealed an age-dependent decrease in PDGF receptor expression, resulting in attenuated proliferative potential of β-cells. In particular, selective inactivation of PDGFRA in β-cells exacerbates sensitivity to the β-cell toxin streptozotocin^41^. Consistent with this, our study showed that *PDGFRA* and *PDGFRB* were upregulated in β-cells from both EC-iFOXO1-KO:STZ/HFD and EC-iDKO:STZ/HFD mice compared with WT:STZ/HFD mice **(Extended Data Fig. 3d)**.

Next, using full-length scRNA-seq data, we investigated the proportions of the 16 clusters in **Fig. 2b**. In agreement with previous studies^1,3^, the proportion of ECs in the pancreas of EC-iFOXO1-KO:STZ/HFD mice was significantly increased by 54% compared with WT:STZ/HFD mice (**Fig. 2c**, **Extended Data Fig. 3a**). Notably, the proportion of β-cells in the total pancreatic cell population was 40% higher in EC-iFOXO1-KO:STZ/HFD mice than in WT:STZ/HFD mice (**Fig. 2c**, **Extended Data Fig. 3a**). These results indicate that EC-specific knockout of FOXO1 increases both ECs and β-cells proportions in diabetes.

### FOXO1 regulates IGF2 expression through CTCF

Methylation of CCCTC-binding factor (CTCF) is crucial for regulation of IGF2 expression^18,19^. Reduced CTCF expression controls IGF2 imprinting, which has been explored in cancer and aging research^20,21^. IGF2 polymorphisms lead to a switch from monoallelic to biallelic imprinting during aging, with a 10-fold increase in expression. This shift is based on a 2-fold reduction in CTCF binding in the intergenic IGF2-H19 region. Downregulation of *CTCF* using small interfering RNA in imprinted prostate cells increases IGF2 expression and relaxes imprinting^20^. However, the boundaries of CTCF dependence controlling IGF2 imprinted expression in diabetes remain unexplored^21^.

To dissect the regulatory role of CTCF on IGF2 expreesion in diabetes, we first examined CTCF expression in ECs using scRNA-seq. Differentially expressed genes (DEG) analysis revealed that *CTCF* was significantly downregulated in pancreatic ECs of WT:STZ/HFD mice compared with WT mice. However, under diabetic conditions, EC-specific knockdown of FOXO1 resulted in upregulation of *CTCF* expression in pancreatic ECs (**Fig. 2d**, **Extended Data Fig. 3e, f**). This suggests that FOXO1 may mediate the regulation of *IGF2* expression through CTCF.

To validate the scRNA-seq observations, we performed western blot and quantitative RT-PCR analysis to further confirm the role of FOXO1 in mediating IGF2 expression by regulating CTCF. Notably, in skin ECs isolated from EC-iFOXO1-KO:STZ/HFD and EC-iDKO:STZ/HFD mice, we observed a significant increase in CTCF expression and a significant decrease in IGF2 expression compared with skin ECs isolated from WT:STZ/HFD mice (**Extended Data Fig. 4c)**. Similarly, *CTCF* expression was significantly upregulated in pancreatic ECs isolated from EC-iFOXO1-KO:STZ/HFD and EC-iDKO:STZ/HFD mice compared with pancreatic ECs isolated from WT:STZ/HFD mice. In contrast, *Igf2* was significantly downregulated in pancreatic ECs from EC-iFOXO1-KO:STZ/HFD, EC-iDKO:STZ/HFD, and EC-iULK1-KO:STZ/HFD mice compared with pancreatic ECs from WT:STZ/HFD mice (**Extended Data Fig. 4f-h)**. These findings imply upregulation of CTCF and downregulation of IGF2 at the protein and transcriptional levels in the context of EC-specific knockout of FOXO1, epsins or ULK1 under diabetic conditions.

At the systemic level, we found that CTCF was significantly reduced in the plasma of WT:STZ/HFD mice compared with WT mice (**Fig. 2e**, **Extended Data Fig. 4d)**. However, compared with WT mice, CTCF levels in the plasma of EC-iFOXO1-KO:STZ/HFD mice were maintained at similar levels, while CTCF levels in the plasma of EC-iDKO:STZ/HFD and EC-iULK1-KO mice were slightly decreased (**Fig. 2e**). Notably, IGF2 levels were significantly elevated in the plasma of WT:SZT/HFD mice compared with WT mice (**Fig. 2e**, **Extended Data Fig. 4d)**. However, IGF2 levels were significantly reduced in the plasma of EC-iFOXO1-KO:STZ/HFD, EC-iDKO:STZ/HFD, and EC-iULK1-KO:STZ/HFD mice compared with WT:STZ/HFD mice (**Fig. 2e**). These results were also recapitulated in transgenic T2D *db/db* mice (**Fig. 2e**, **Extended Data Fig. 4e, i, j)**. Compared with WT:*db/db* mice, CTCF levels in the plasma of EC-iDKO:*db/db* mice were significantly increased, while IGF2 levels were significantly decreased (**Fig. 2e**, **Extended Data Fig. 4e)**. Notably, the levels of IGF2R, an IGF2 scavenger, were increased in EC-iFOXO-KO:STZ/HFD mice compared with WT:SZT/HFD mice (**Extended Data Fig. 4h)**.

In summary, our findings reveal that EC-specific knockout of FOXO1 can increase CTCF expression and reduce IGF2 levels under diabetic conditions, indicating that FOXO1 regulates IGF2 expression through CTCF.

### Depletion of EC-FOXO1 depletion enhances the activity of IGF2R interaction in the diabetic islet niche

Next, we further evaluated how pancratic ECs impact β-cells through the IGF2-IGF2R system in diabetes. Notably, *IGF2R* was upregulated in ECs of EC-iFOXO1-KO:STZ/HFD mice compared with WT:STZ/HFD mice. In contrast, *IGF2* was downregulated in ECs of EC-iFOXO1-KO:STZ/HFD mice compared with WT:STZ/HFD mice (**Fig. 2f**). Except for ECs, *IGF2* and *IGF2R* expression levels in β-cells exhibit similar manners. These results suggest that the IGF2-IGF2R system is pivotal in mediating communication between pancreatic ECs and β-cells, potentially affecting the development of diabetes. To explore this further, we performed RNA-seq analysis to assess the expression of IGF2R. Notably, EC-iFOXO1-KO:STZ/HFD and EC-iDKO:STZ/HFD mice showed upregulation of *IGF2R* in EC compared with WT:STZ/HFD mice (**Fig. 2g**, **Extended Data Fig. 5a, g**).

To dissect how ECs communicate with endocrine cells, specifically to evaluate how EC-specific knockout of FOXO1 can modulate gene expression of β-cells in the diabetic pancreatic islet niche, we utilized NicheNet^42^ to identify crucial ligands and receptors derived from ECs and endocrine cells and predicted their associations. Our primary objective was to understand how genes differentially expressed in ECs affect β-cell function under diabetic conditions. Notably, we observed that, in addition to other key factors such as *INS1, INS2, IGF1*, and *IGF1R,* the association between ECs and β-cells prominently involved the expression of *IGF2R* in the EC- iFOXO1-KO:STZ/HFD compared with the WT:STZ/HFD, suggesting that IGF2R may function as a scavenger to inhibit and eliminate excess IGF2 in the diabetic pancreatic islet niche (**Fig. 2h**). Our results demonstrate that EC-specific knockout of FOXO1 significantly increases IGF2R interactive activity in the diabetic islet niche, which may play an important role in regulating β-cell function in diabetes.

### IGF2R promotes α-to-β cell conversion in EC-FOXO1-depleted diabetic mice

Previous studies reported that adult pancreatic exocrine cells can be reprogrammed into β-cells^28,32^. Further lineage tracing analyses revealed the conversion of adult pancreatic α-cells to β-cells following the massive loss of β-cells in diabetes^28^. The transcription factors Pdx1, Mafa, and Nkx6-1 can reprogram adult mouse pancreatic exocrine cells into β-cell-like cells^29,32^. Human islets also exhibit this plasticity, notably switching of cellular identity that may occur under diabetic conditions^32^. To investigate the involvement of these transcription factors in endocrine cell plasticity within our diabetic STZ/HFD models, we assessed the expression of transcription factors including Mafa, Pdx1, Nkx6-1, Gata6, Foxa2, Neurod1, Pax6, Mafb, Nkx2-2, Isl1, Sox9, Ptf1a, and Gata4 in pancreatic cells of EC-iFOXO1-KO:STZ/HFD and WT:STZ/HFD mice through scRNA-seq analysis. Notably, we identified significantly elevated expression levels of *Mafa, Sox9, and Gata6* in ECs, β-cells, α-cells, δ-cells, γ-cells, acinar cells, ductal cells, and acinar ductal metaplasia (ADM) cells from EC-iFOXO1-KO:STZ/HFD mice compared to cells from WT:STZ/HFD mice. *Pdx1, Nkx6-1, and Foxa2* were significantly upregulated in β-cells and/or acinar cells, ductal cells, and ADM cells of EC-iFOXO1-KO:STZ/HFD mice compared with WT:STZ/HFD mice (**Extended Data Fig. 4a**). These findings suggest that Mafa, Sox9, and Gata6 may play key roles in islet endocrine cell plasticity under diabetes.

We further evaluated scRNA-seq data from WT:STZ/HFD and EC-iFOXO1-KO:STZ/HFD mice to study the reprogramming process of pancreatic endocrine cells, especially α-to-β cells, at single-cell resolution. Using Uniform Manifold Approximation and Projection (UMAP) visualization, we identified two distinct cell populations of α-cells and β-cells (**Fig. 2i, left**). The pseudotemporal ordering algorithm was employed to reconstruct the cellular gene expression profiles and developmental trajectory without relying on prior knowledge of genes defining progression. This analysis revealed a main trajectory with a few minor branches, allowing the classification of three pseudotime-dependent progression states for α-to-β cells: "early," "mid," and "late" (**Fig. 2i, right**). The "early" and "mid" stages are upregulated α-cell gene markers (*Gcg*), while the "late" stage is upregulation of β-cell gene markers (*Ins1 and Ins2*). Significantly, *Ins1 and Ins2* were upregulated during the α-to-β cell progression states in the EC-iFOXO-KO:STZ/HFD mice compared to that of WT:STZ/HFD, suggesting that EC-specific knockout of FOXO1 can increase α-to-β cell progression to compensate for β-cells loss under diabetic conditions (**Fig. 2j**). Notably, although *Igf1* and *Igf1r* exhibited similar expression patterns, the proportion of cells expressing *Igf2r* was observed to increase during α-to-β cell progression. Specifically, the proportion of cells expressing *Igf2r* was significantly higher in β-cells compared with α-cells. Furthermore, *Igf2r*-expressing cells (14.31%) were more abundant in β-cells from EC-iFOXO1-KO:STZ/HFD mice than in β-cells from WT:STZ/HFD mice (8.67%) (**Fig. 2j**, **Extended Data Fig. 4b**). These findings suggest that IGF2R plays a critical role in the conversion of α-to-β cells, and that EC-specific depletion of FOXO1 promotes this conversion under diabetic conditions.

### EC-FOXO1 regulates β-cell fate in diabetes via the IGF2-IGF2R system

KEGG pathways analysis revealed that EC-iFOXO1-KO:STZ/HFD elicited many differentially expressed genes related to vascular development, angiogenesis, insulin signaling, autophagy, and wound healing pathways compared with WT:STZ/HFD (**Extended Data Fig. 5b-f, h, i**). Given that upregulation of IGF2R and downregulation of IGF2 play a key role in improving diabetes in EC-specific knockout FOXO1 mice, we further investigated in depth the mechanism by which FOXO1 regulates β-cell function and fate in diabetes.

Excess unbound soluble IGF2 destroys β-cells in the pancreas and is a major cause to the pathology of diabetes^13–15,26^. We first investigated whether IGF2 could induce β-cell apoptosis. Notably, the treatment of β-TC-6 cells with IGF2 *in vitro* resulted in more pronounced β-cells loss in a dose-dependent manner, with approximately 47.4% of β-cells showing late apoptosis after overnight incubation with an IGF2 concentration of 3 pg/mL (**Fig. 3a, b**, **Extended Data Fig. 6a, b**).

EC-specific knockout of FOXO1, epsins, or UlK1 increased IGF2R expression (**Fig. 1j-o**, **Fig. 2f, g**, **Extended Data Fig. 2, 4h, i**) and decreased IGF2 expression (**Fig. 1i**, **Fig. 2e, f**, **Extended Data Fig. 4c-e, g**). Increased IGF2R is predicted to reduce IGF2 and restore pancreatic β-cell loss in diabetes. Therefore, we sought to investigate whether these ECs affect viability of β-cells. Of note, ECs isolated from EC-iFOXO1-KO:STZ/HFD, EC-iDKO:STZ/HFD, and EC-iULK1-KO:STZ/HFD mouse pancreas significantly reduced early apoptosis of β-cells compared to WT:STZ/HFD (**Extended Data Fig. 6c-e**).

We co-cultured β-cells with ECs isolated from WT:STZ/HFD, EC-iFOXO1-KO:STZ/HFD, EC-iDKO:STZ/HFD, and EC-iULK1-KO:STZ/HFD mouse pancreas, and added appropriate doses of IGF2 (1.5 pg/mL, approximately 2 times the plasma IGF2 level in WT:STZ/HFD, as shown in **Fig. 1i**) to study β-cells loss (**Fig. 3c**). Notably, β-cell mortality induced by IGF2 was reduced by 87.5% and 72.3% when co-cultured with pancreatic ECs from EC-iFOXO1-KO:STZ/HFD and EC-iDKO:STZ/HFD mice, respectively, compared to that of WT:STZ/HFD mice (**Fig. 3d-f**). The later apoptosis ratio of β-cells induced by IGF2, when co-cultured with the pancreatic ECs from EC-iFOXO1-KO:STZ/HFD, EC-iDKO:STZ/HFD, EC-iULK1-KO:STZ/HFD mice, was significantly decreased by 71.3%, 73.5%, and 51.5%, respectively, compared to that of WT:STZ/HFD mice, indicating that EC-specific knockout of FOXO1, as well as epsins, or ULK1 prevents β-cell apoptosis through the mediated IGF2-IGF2R system (**Fig. 3d, e, g**)

In brief, our findings suggest that the exogenous addition of IGF2 to β-cells *in vitro* promotes massive β-cell death. In diabetes, EC-specific deficiency of FOXO1, as well as epsins and ULK1, respectively increases β-cell survival through upregulation of IGF2R and inhibition of IGF2.

### FOXO1 regulates epsin-dependent IGF2R endocytosis in diabetes

Gene concept network analysis by scRNA-seq revealed that compared to the WT:STZ/HFD control, EC-FOXO1 depletion in diabetes elicited many differentially expressed genes, including *IGF2R, CTCF, Vegfa, Kdr, Epn2, Ins1, Ins2, Igf1, Pdgfra,* and *Map1lc3b*. These highly

expressed genes are mainly involved in insulin-like growth factor binding, regulation of gene expression by genomic imprinting, insulin signaling, IGF signaling, and VEGF signaling (**Fig. 3h**).

Previous studies have shown IGF2R facilitates the trafficking of lysosomal enzymes from the trans-Golgi network to endosomes and then to lysosomes^17,43,44^. Consistently, our colocalization studies showed that IGF2R was co-localized with the autophagy marker LC3B and the lysosome marker LAMP1 in pancreatic ECs from WT:STZ/HFD, implying that IGF2R is capable of degrading and clearing excess free IGF2 in diabetes through the endocytic pathway (**Extended Data Fig. 6f**).

Given the primary role of IGF2R as a scavenger receptor, binding IGF2 at the cell surface and promoting its internalization into lysosomes for degradation, we investigated whether epsins, a family of endocytic adaptor proteins that select specific cargo for internalization by endocytosis^5–7,24^, might contribute to IGF2R internalization. We conducted a co-immunoprecipitation to assess the physical interaction between epsins and IGF2R. Notably, our results revealed that the full-length epsin 1, the N-terminal homology (ENTH) domain, and/or the ubiquitin-interacting motif (UIM) of epsin 1 exhibited interaction with IGF2R. Conversely, no interaction was observed between the C-terminal domain of epsin 1 and IGF2R (**Fig. 3i**). Consistently, Epsin 1 co-localizes with IGF2R in the cytoplasmic region of pancreatic ECs isolated from WT:STZ/HFD mice (**Extended Data Fig. 6g**). These findings suggest a direct interaction between epsin 1 and IGF2R.

Hence, considering the observed FOXO1-mediated upregulation epsins (**Fig. 1a**), our data suggest that FOXO1 can regulate epsin-dependent endocytosis of IGF2R in a diabetic context.

### EC-FOXO1 depletion enhances angiogenesis by inhibiting autophagosome formation

Given that FOXO1 upregulates epsin 1 and ULK1 (**Fig. 1a–d**, **Extended Data Fig. 1a–d, f**), we hypothesized that FOXO1 could modulate the association of epsin 1 with LC3B and ULK1, leading to the formation of autophagosomes together with VEGFR2 in diabetes (**Fig. 4a**, **Extended Data Fig. 7e)**. The formation of these autophagosomes promotes VEGFR2 degradation, thereby impairing angiogenesis in diabetes^6,38^.

We first sought to explore the potential synergistic effects of EC-specific defects in FOXO1, epsins, and ULK1 on angiogenesis and their potential impact on increasing IGF2R levels, which may contribute to diabetes protection. Leveraging electron microscopy (EM), we quantitatively analyzed autophagosomes, which were characterized by electron-dense vacuoles with double membranes, in pancreatic ECs isolated from STZ/HFD and *db/db* mice. Autophagosomes exhibited a significant reduction in pancreatic ECs isolated from EC-iFOXO1-KO:STZ/HFD and EC-iDKO:STZ/HFD mice when compared to WT:STZ/HFD mice (**Fig. 4b-e**). Notably, a significant increase was observed in ECs obtained from WT:*db/db* mice compared to those from iDKO:db/db mice, while autophagosomes were significantly reduced in ECs isolated from WT:*db/db* mice treated with ULK1 siRNA compared to control siRNA (**Extended Data Fig. 7a-d**). Our results indicate that EC-specific knockout FOXO1, epsins or knockdown UlK1 can reduce autophagosome formation in diabetes.

Interestingly, the intracellular kinase domain of VEGFR2 (amino acids 996-1004) contains the LC3B-interacting region (LIR) motif that plays a critical role in LC3B binding^45^ (**Extended Data Fig. 7e**). Co-immunoprecipitation analysis revealed that LC3B interacts with VEGFR2 through this LIR motif (**Extended Data Fig. 7f, g**). Notably, immunofluorescence analysis demonstrated co-localization of epsin 1, LC3B and VEGFR2 in HUVECs treated with 50 μM MGO (**Fig. 4f**). Furthermore, in co-immunoprecipitation assays, methylglyoxal (MGO)-treated HUVECs exhibited enhanced interactions of epsins with endogenous ULK1 and LC3B (**Extended Data Fig. 7h**). This interaction was further confirmed in HEK 293T cells expressing full-length or domain-deleted FLAG-epsin 1 constructs in pcDNA3 vectors, which were co-transfected with LC3B or ULK1 expression vectors (**Extended Data Fig. 7i, j**). Our findings suggest that full-length epsin 1 and/or ENTH and UIM domains of epsin 1 can interact with LC3B and ULK1, respectively. In the context of diabetes, epsin 1 can bind to VEGFR2 via LC3B to form an autophagosome complex, where VEGFR2 will be selectively degraded through LIR-mediated interactions (**Fig. 4 a, f**, **Extended Data Fig. 7e-g, h-j**).

To gain a deeper insight into the impact of resultant autophagosomes on angiogenesis in the context of diabetes, we conducted a corneal micropocket assay. These analysis revealed significantly increased vessel density, branch points, and filopodia projections in EC-iFOXO1-KO:STZ/HFD, EC-iDKO:STZ/HFD, and EC-iULK1-KO:STZ/HFD compared with those in WT:STZ/HFD mice, upon implantation with VEGF-A-containing micropellet (**Fig. 4g–i**). This suggests that EC-specific depletion of FOXO1, epsins, or ULK1 restores angiogenesis in diabetes. To validate the *in vivo* findings, we examined the impact of EC-FOXO1 depletion on angiogenesis *in vitro*. Consistent with *in vivo* observations, ECs from EC-iFOXO1-KO:STZ/HFD mice exhibited a significant increase in tube formation (**Extended Data Fig. 8a, b**), proliferation (**Extended Data Fig. 8c, d**), and migration (**Extended Data Fig. 8e, f**) in responses to VEGF-A compared to ECs from WT:STZ/HFD mice. ECs from EC-iDKO:STZ/HFD and EC-iULK1-KO:STZ/HFD displayed similar behavior *in vitro* (Data not shown).

Overall, our findings reveal that the specific knockout of FOXO1, epsins, or ULK1 in ECs can enhance angiogenesis by inhibiting autophagosome formation in diabetes.

### EC-FOXO depletion accelerates diabetic wound healing

Lastly, we sought to investigate whether EC deficiency of FOXO1, as well as epsins or ULK1 could improve wound healing under diabetes.

Notably, the dermal wounds induced by biopsy punch in EC-iFOXO1-KO:STZ/HFD, EC-iDKO:STZ/HFD, and EC-iULK1-KO:STZ/HFD mice remarkably accelerated the wound healing process, leading to faster closure of the initial wounds compared to that observed in WT:STZ/HFD mice (**Fig. 4j, k**). While the wound closure in EC-iDKO:*db/db* mice was delayed compared to normal WT mice at day 12, it exhibited significant improvement and acceleration when compared to WT:*db/db* mice at day 24 (**Extended Data Fig. 10a, b**). Histological examination of wounded skin tissues revealed elevated IGF2 levels in WT:STZ/HFD mice as compared to EC-iFOXO1-KO:STZ/HFD mice (**Extended Data Fig. 9a-c**). IGF2 levels were also assessed in the pancreas of these mice. Immunofluorescence analysis showed that IGF2 levels were significanlty reduced in pancreatic tissues from EC-iFOXO1-KO:STZ/HFD, EC-iDKO:STZ/HFD, and EC-iULK1-KO:STZ/HFD mice compared with WT:STZ/HFD mice (**Extended Data Fig. 9d-f**). Furthermore, the expression level of IGF2R was significantly increased in ECs isolated from the skins of EC-iFOXO1-KO:STZ/HFD, EC-iDKO:STZ/HFD, and EC-iULK1-KO:STZ/HFD mice compared with ECs isolated from the skins of WT:STZ/HFD mice (**Fig. 4l, m**). Collectively, our results suggest that the EC-specific depletion of FOXO1, as well as epsins, or ULK1 accelerates the healing of diabetic wounds, and the IGF2-IGF2R system may play a critical role in the wound healing process.

In conclusion, FOXO1 functions as a repressor of CTCF, leading to increased IGF2 expression in diabetes. Meanwhile, FOXO1 upregulates epsins and ULK1 to promote the degradation of IGF2R and VEGFR2, ultimately inhibiting angiogenesis through autophagosome formation. Thus, excessive IGF2 levels and reduced angiogenesis impair β-cell function and survival, exacerbate diabetes, and impede the wound healing process (**Extended Data Fig. 10c**). However, EC-FOXO1 deficiency results in elevated CTCF levels and effectively blocks IGF2 expression in diabetes. Depletion of EC-FOXO1 also prevents autophagosome formation and lysosomal degradation of IGF2R and VEGFR2, thereby increasing angiogenesis. Therefore, reducing IGF2 levels and enhancing angiogenesis are expected to improve β-cell function and survival, ameliorate diabetes, and accelerate skin wound healing (**Extended Data Fig. 10d**).

## Discussion

Here, we provide evidence for the presence of excess IGF2 in the pancreas of diabetic patients, and in the pancreas, skin, and plasma of STZ/HFD-induced diabetic and *db/db* mice. Excess IGF2 in diabetic mice leads to β-cell dysfunction and death and may be involved in diabetic wound healing^12–15,25^. Importantly, we observed, for the first time, that EC-specific depletion of FOXO1 significantly increase both IGF2R and CTCF levels while notably decreasing IGF2 level in the context of diabetes (**Fig. 1**, **Fig. 2**, **Extended Data Fig. 1**-**6**). IGF2 can induce β-cell apoptosis in a dose-dependent manner. EC-FOXO1 depletion, as well as epsins and ULK1 can prevent IGF2-induced β-cell dysfunction and loss (**Fig. 3a-g**, **Extended Data Fig. 1k-m**, **Extended Data Fig. 6a-e**).

IGF2 plasma levels are notably higher than in postnatal life across species from sheep, guinea pigs to man^46,47^, whereas postnatal levels in rats and mice were barely detectable^46^. However, these studies were largely conducted under normal physiological growth and metabolism conditions^15,16,48^ and the observed outcomes may be influenced by phylogenic placement^49,50^. Human endocrine pancreatic single-cell transcriptomics revealed significantly increased IGF2 expression in β-cells from T2D patients^33^. Studies have demonstrated that the methylation of a CTCF-dependent boundary regulates the imprinted expression of the *Igf2* gene^18–21^. Consistently, our scRNA-seq transcriptome profiling and bulk RNA-seq analysis reveal that *CTCF* is downregulated in WT:STZ/HFD mice compared to WT mice, however, EC-specific knockdown of *FOXO1* leads to upregulation of *CTCF* expression under diabetic conditions, implying *FOXO1* mediates *Igf2* expression through *CTCF* regulation (**Fig. 2d, g**, **Extended Data Fig. 3e, f**). Furthermore, immunofluorescence and western blotting studies demonstrated that FOXO1 regulates IGF2 expression through CTCF (**Fig. 2e**, **Extended Data Fig 4c-i**). Previous studies have shown that IGF2R functions as a scavenger, promoting the internalization and degradation of IGF2^15–17,27^. Moreover, methylation of the CTCF-dependent boundary governs IGF2 expression^18–21^. These dual pathways, involving the suppression of IGF2 levels by IGF2R^8,15^ and CTCF^18–21^, respectively, are highly consistent with our discovery that EC-FOXO1 depletion effectively diminishes elevated IGF2 levels in diabetes. This reduction of IGF2 has the potential to improve β-cell dysfunction and loss, and ultimately ameliorate diabetes. The precise mechanism by which FOXO1 negatively regulates CTCF levels in diabetes warrants further investigation, leveraging comprehensive 3D chromatin transcriptional profiling, such as ATAC-seq.

Single-cell transcriptomic profiles of the mouse pancreas identified significantly elevated expression levels of *Mafa, Sox9, and Gata6* in various cell types, including pancreatic ECs and endocrine cells from EC-iFOXO1-KO:STZ/HFD mice compared to cells from WT:STZ/HFD mice (**Extended Data Fig. 4a**). Trajectory analysis indicates that IGF2R plays a crucial role in the conversion of α-to-β cells, and the depletion of EC-FOXO1 promotes this conversion under diabetic conditions (**Fig. 2i, j**, **Extended Data Fig. 4b**). However, the precise molecular mechanisms and clinical applications of transcription factors such as *FOXO1, Mafa*, *Sox9*, *Nkx6-1*, and *Foxa2* in mediating the plasticity of pancreatic endocrine cells, as well as the role of IGF2-IGF2R system in this process remain to be further investigated.

Previous studies link EC FOXO1 involvement in the development of diabetic vascular complications in patients^51–55^, however molecular mechanisms remain elusive. Our studies here established that endothelial FOXO1 governs epsin-dependent angiogenesis via the autophagosomal-lysosomal degradation of VEGFR2. VEGFR2 is a key angiogenic factor required for wound healing but it is markedly decreased in the endothelium of diabetic patients ^6,7,24,56–58^. In diabetic mouse models, EC-specific knockout or knockdown of FOXO1 inhibits epsins and the autophagosome assembly protein ULK1. This inhibition prevents VEGFR2 degradation, promotes angiogenesis, and ultimately facilitates wound healing in diabetes.

To summarize, our current study represents a novel therapeutic avenue for reducing β-cell dysfunction and loss, ameliorating diabetes, and promoting wound healing.

### Methods Mouse Models

All *in vivo* experiments on animals were approved by the Institutional Animal Care and Use Committee at Boston Children’s Hospital, Harvard Medical School, Boston, MA. The endothelial cell (EC) specific deletion of epsins (*Epn1*^fl/fl^, *Epn2*^−/−^) mice (EC-iDKO) have been previously described^59,60^. Mice harboring floxed alleles of *FoxO1* (*FoxO1*^fl/fl^) were obtained from Sudha B. Biddinger and C. Ronald Kahn and have been described previously^35^. The conditional deletion of *FoxO1* mice was generated by breeding these mice with EC-specific Cre transgenic mice (iCDH5-Cre ERT2) to create EC-iFOXO1-KO mice. To generate an endothelial-specific deletion of the Ulk1 mouse strain (EC-iUlk1-KO), *Ulk1*^fl/fl^ mice were crossed with Tek-CreTg transgenic mice (both from the Jackson Laboratory; Bar Harbor, ME). *FoxO1*^fl/fl^; *Epn1^fl/fl^,Epn2^-/-^*; *Ulk1*^fl/fl^; or *Epn1^fl/fl^,Epn2^-/-^, Ulk1*^fl/fl^ littermate mice without iCDH5-Cre-ERT2 or wild type (WT) mice with iCDH5-Cre-ERT2 were utilized as controls and denoted as WT. For Cre recombinase activation, EC-iFOXO1-KO, EC-iDKO, and EC-ULK1-KO or EC-iTKO mice were administered 4-hydroxytamoxifen (Hello Bio, HB6040-50mg, dissolved in a 9:1 mixture of DMSO and ethanol, 5–10 mg/kg body weight) 7 times every other day, starting at 8∼10 weeks old. Mice were rendered diabetic through intraperitoneal injection of a low-dose streptozotocin (STZ, Sigma-Aldrich, S0130-500MG, 50 mg/kg) regimen, as previously described^7^. Hyperglycemia was defined as a fasting (6 h) blood glucose level of > 200 mg/dL for > 1 week after STZ administration. Mice were subsequently fed a high fat diet (HFD, 60 kcal% fat, Research Diets Inc., D12492) and denoted as WT:STZ/HFD (WSH), EC-iDKO:STZ/HFD (DSH), EC-iFOXO1-KO:STZ/HFD (FSH), and EC-iULK1-KO:STZ/HFD (USH), or EC-iTKO:STZ/HFD (Triple Knockout, TSH).

Leptin receptor-deficient diabetic *db/db* mice were purchased from Charles River (JAXTM mice). EC-iFOXO1-KO and EC-iDKO mice were crossed with *Lepr^-/-^ db/db* mice respectively, to generate the endothelial-specific deletion of the FOXO1 or epsins in the context of diabetic *db/db* mice (EC-iFOXO1-KO:*db/db* or EC-iDKO:*db/db*). The EC-iDKO:*db/db* mice (*Lepr^-/-^ db/db*) and EC-iFOXO1-KO:*db/db* (*Lepr^+/-^ db/db*) mice were obtained. However, we encountered challenges in obtaining viable EC-iFOXO1-KO:*db/db* (*Lepr^-/-^ db/db*) mice, possibly due to the embryonic lethality during the breeding process.

Determination of sample size was guided by the results of our preliminary experiments. All STZ/HFD and *db/db* mouse models were selected according to their correct genotypes. Mice were housed in well-ventilated cages with constant temperature and humidity (20-23°C, 50-60%) in a pathogen-free controlled environment with a 12-h light-dark cycle set with light on from 06:00 to 18:00. For genotyping of *db/db* mice, the primers were from Jackon laboratory: forward, 5’-AGAACGGACACTCTTTGAAGTCTC-3’ (oIMR0985), reverse, 5’-CATTCAAACCATAGTTTAGGTTTGTGT-3’ (oIMR0986).

### Human samples

Studies using human samples were performed according to approved human research guidelines at Boston Children’s Hospital, Harvard Medical School, Boston, MA. Human pancreas paraffin tissue blocks from human adult healthy control and diabetic patients were purchased from BioChain Insititute, Inc. The paraffin-embedded pancreatic sections, were deparaffinized and rehydrated using xylene, followed by washing with a descending series of ethanol (100%, 95%, 70%, 50%), and water. Deparaffinized slides were subjected to antigen retrieval using Retrievagen A (pH 6.0, BD Bioscience) following to the manufacturer’s protocol. The antigen-retrieval slides were then processed for immunofluorescence staining described below.

### Chromatin immunoprecipitation (ChIP) assay

Within -20 kb upstream of the mouse *Epn1* transcription start site (TSS), there are approximately nine fragments containing the FOXO1 binding sequence element 5’-TGTTT-3’. Likewise, 6 fragments within −20 kb upstream of the mouse *ULK1* TSS also contain this element. PCR primers for chromatin immunoprecipitation sequencing (ChIP-Seq) were designed to target 9 fragments of *Epn1* and 6 fragments of *ULK1*. ChIP-Seq assay was performed to validate FOXO1-binding fragments using a ChIP assay kit (QIAGEN) following the manufacturer’s protocol and previously described methods^7^. Briefly, ECs isolated from WT:STZ/HFD mice were cultured in 10-cm dishes to 70% confluence, cross-linked with 1% formaldehyde for 10 min at 37°C, and then lysed. DNA was sheared (500-1000 bp) using sonication and preclear lysates were precleared with DNA-Protein A Agarose. Immunoprecipitation was performed overnight at 4°C using anti-FOXO1(Cell signaling), anti-IgG (QIAGEN kit), and anti-GAPDH (QIAGEN kit) antibodies. Immune complexes are collected, eluted, and cross-link reversal and DNA recovery are performed. Purified DNA served as template for PCR amplification of FOXO1 binding sites. ChIP-seq PCR shows that the significant strong binding signal located in the frist fragment within the promoter region of both *Epn1* and *ULK1*. The primer pairs for these fragments: *Epn1* (Forward, 5’-GCCTCTTTTGCCATGACTCCTG-3’, Reverse, 5’-GCCCAGTAGTGGTGTCAAGGGG-3’); *ULK1* (Forward, 5’-CGCCTAAGAGGTCTTGAGCTCC-3’, Reverse, 5’-CCCGGCGCTGACAGTCGGGACC-3’).

### Luciferase Assay

The promoter fragments of *Epn1* and *ULK1* contain FOXO1 binding sequence elements: 5’-gTGTTTcc-3’ is located approximately 110 bp upstream of the *Epn1* transcription start site (TSS), and 5’-tTGTTTcc-3’ is located 324 bp upstream of *ULK1* TSS. These sequences are consistent with previously identified FOXO1 binding sites^3^. These two fragments which included FOXO1 binding sequences were inserted into the luciferase reporter vector pGL4.23 [luc2/minP] (Promega) in the Forward orientation. Primary ECs isolated from WT:STZ/HFD mice were transfected with 1 μg of reporter plasmid and 1 μg of effector plasmids using Lipofectamine® LTX Plus DNA transfection reagents, following the manufacturer’s protocol (ThermoFisher Scientific). After 48 hours of transfection, cells were harvested, and cell extracts were assayed for luciferase activity using a Dual Luciferase Reporter Assay System (Promega). Renilla luciferase plasmid (Promega) was co-transfected as a control to normalize transcription efficiency. Mutations were introduced into the FOXO1 binding sites within the *Epn1* or *ULK1* promoter-driven luciferase reporter plasmids using the Q5^®^ Site-Directed Mutagenesis Kit (New England BioLabs, Inc., E0554S), according to the manufacturer’s instructions. The primer pairs for the Luciferase constructs and mutants of the FOXO1 binding sequences: pGL4.23-Epn1_site1 (Forward, 5’-gcgcGCTAGCTTTAAGACAGAAGTTAACGGG - 3’; Reverse, 5’- GCGctcgagATTGTGCCCAGTAGTGGTGTCAA - 3’); mutant primers of pGL4.23- Epn1_Site1_Mut (Forward, 5’ - tgctaaccccccaaacccCCCCAGATGTGGTGATCTAC -3’, Reverse: 5’- gcgcgaaagagatgtggagGCAACCAGCACCCAAGCT -3’). pGL4.23-ULK1_site1 (Forward, 5’-GCGgctaGCCAGCCAGTTCTACTGAGTGAG-3’, Reverse, 5’- GCGctcgAGCGCGCGCGGGCCGGGGGCGCG-3’); mutant primers of pGL4.23- ULK1_Site1_Mut (Forward, 5’- tctcccTGCGGGGCTCCGAGCCGC – 3’, Reverse: 5’-tagaagATCCGCGGCACCCGGGCC – 3’).

### GTT and ITT

Glucose tolerance test (GTT) and insulin tolerance test (ITT) were conducted as previously described^7^. For GTT, mice were fasted for 16 hours and received an Intraperitoneal (I.P.) administration of a calculated 10% D-glucose (Sigma-Aldrich, G8270) dose at a ratio of 2.5 g/kg body weight. For ITT, mice were fasted for about 4-6 hours and received an I.P. injection of insulin (Humulin R, Lilly) at a calculated dose of 0.75 U/kg body weight. Insulin was dissolved in a 0.9% sterile saline solution. Blood glucose levels in the GTT and ITT were measured at 0, 15, 30, 60, 90, and 120 minutes after D-glucose or insulin injection using tail vein samples and a glucometer (CVS Health).

### GSIS assay

Pancreas islets from STZ/HFD and *db/db* mice were isolated after collagenase perfusion (collagenase XI, Sigma, C7657), pancreatic digestion, and purification as previously described with modifications^61^. Equal amounts of handpicked islets from each mouse were cultured in 6-well plates in RPMI 1640 medium supplemented with 10% (v/v) fetal bovine serum (Neuromics, FBS007) and 1% penicillin-streptomycin. (ThermoFisher Scientific, 15140122), with 1 ml of this medium overnight. After 24 h of incubation, the culture medium was harvested, centrifuged at 4°C and 12,000 r.p.m. for 5 minutes, and approximately 100 μL of the supernatant was saved as the initial baseline level of insulin secretion by the islets. The islets were then transferred to the same fresh RPMI 1640 medium containing 25mM glucose and continually incubated in a 37°C incubator. After 2 h of stimulation, the culture mediums were collected, centrifuged at 4°C and 12,000 r.p.m. for 5 m, and approximately 100 μL of the supernatants were retained and stored at -80°C for insulin measurement. Insulin levels were measured using the Ultra-Sensitive Mouse Insulin ELISA kit (Crystal Chem, 90080), as described below.

### Plasma collection

Blood collection from mice was performed as previously described^62^. Briefly, the mice were restrained in a 50-ml tube with some air holes at the bottom (BD Biosciences, Heidelberg, Germany). Mouse legs were isolated above the knee and the fur was shaved away with a GT420 electric shaver (Braun, Suhl, Germany) until the saphenous vein was visible, after which a 27-gauge sterile needle was used to puncture the lateral saphenous vein. Blood was then collected in the blood collection tube (BD Microtainer^®^) or in the 1.7mL microcentrifuge tube containing 50 μL 0.5M EDTA using the microhematocrit heparinized capillary tube (Fisher scientific), and bleeding was stopped with a cotton pad. The collected blood was centrifuged at 1,500 r.p.m. at room temperature for 8 min. The supernatant was further centrifuged at 12,000 r.p.m. for 10 min at 4^0^C to obtain clear plasma for western blot assay, as well as analysis of plasma cholesterol, triglyceride, insulin, and IGF2 levels.

### Insulin and IGF2 measurement

To validate the presence of insulin and IGF2 in the plasma of WT, WT:STZ/HFD, EC-iFOXO-KO:STZ/HFD, EC-iDKO:STZ/HFD, and EC-iULK1-KO:STZ/HFD mice or *db/db* mice, different batches of plasma from these mice were collected for analysis using the sandwich enzyme-linked immunosorbent assay (ELISA) kit. Plasma insulin and IGF2 were measured using the Ultra-Sensitive Mouse Insulin ELISA kit (Crystal Chem, 90080) and the Mouse IGF-II/IGF2 DuoSet ELISA kit (R&D systems) respectively, following the manufacturer’s instructions. For the insulin ELISA assay, 5 μl of plasma from STZ/HFD and db/db mice was used, with a Wide Range Assay (0.1-12.8 ng/mL). For the IGF2 ELISA assay, 10 μL of plasma was used for a Wide Range Assay (33.3-2000 pg/mL). Signals from the insulin ELISA plates were measured at absorbance 450 and 630 using an Opsys MR^TM^ Microplate reader (Dynex Technologies). The signal of the IGF2 ELISA plate was measured at absorbance 450 and 595 using FilterMax™ F3 and F5 multimode microplate readers (VWR).

### Cholesterol and triglyceride analysis

Total cholesterol and triglycerides were determined in mouse plasma colorimetrically using Ace Axcel® Clinical Chemistry System (Alfa Wassermann, West Caldwell, NJ) as described earlier^63–66^.

### Single-cell RNA-seq of pancreas

WT, WT:STZ/HFD, EC-iDKO:STZ/HFD, EC-iFOXO1-KO:STZ/HFD, and EC-iULK1-KO:STZ/HFD, and *db/db* mice under isoflurane anesthesia were perfused transcardially with about 30 mL of 1X PBS until the livers were cleared of blood. Pancreases were minced into small pieces on ice and then incubated in pre-warmed 10 mL DMEM (ThermoFisher Scientific, 11-966-025) with Collagenase Type II (1mg/mL, ThermoFisher Scientific, 17-101-015), at 37^0^C for 45 min with shaking. To prevent over-digestion cells and prevent them from adhering and aggregating into a ’clumping’ state, digested pancreatic tissues were immediately placed on ice and 3 mL of FBS was added. The digested tissues were then filtered through a 70-μm cell strainer (ThermoFisher Scientific) and followed by centrifugation at 1,200 rpm for 5 min at 4^0^C. Cells were further rinsed with ice-cold DMEM and filtered through a 40-μm cell strainer three times to obtain pure single cells. Trypan blue (ThermoFisher Scientific) was used to count and assess the concentration and viability of single cell suspensions, with all samples containing more than 95% viable cells. Cell pellets were then resuspended in a cell resuspension buffer at 1,000 cells per μL for single-cell RNA-seq (scRNA-seq) library preparation. The rest of cells were used for further flow cytometry analysis, ECs isolation, and RNA isolation, western blot analysis.

The quality control of Post Library Construction and cDNA libraries quality control were performed at the Biopolymers facility, Department of genetics, Harvard Medical School. The sequencing was performed in MedGenome Inc. Reagents for scRNA-seq: Chromium Next GEM Single Cell 3’ Reagent Kits v3.1 (10 X Genomics): GEM Single Cell 3’GEM Kit v3.1(PN-1000123), GEM Single Cell 3’Library Kit v3.1 (PN-1000157), GEM Single Cell 3’Gel Bead Kit v3.1 (PN-1000122), DynabeadsTM MyOneTM SILANE (PN-2000048), GEM Chip G Single Cell Kit, 48rxns (PN-1000120), Single Index Kit T Set A, 96 rxns (PN-1000213), SPRIselect Reagent (Cat# B23317), phorbol 12-myristate 13-acetate (PMA) (Cat#p8313).

### Single-cell RNA-seq analyses

To obtain high-quality scRNA-seq data, we conducted two batches of pancreas scRNA-seq experiments, labeled as P83 and P150-C1.

For the scRNA-seq of P83, raw single-cell RNA sequencing data underwent preprocessing using CellRanger v6.1.1 (10x Genomics), which included aligning reads to mouse reference genome, and generating cellxgene expression count matrices. Subsequently, using the generated expression count matrices, an in-house scRNA-seq pipeline was established based on the Seurat R package (R v3.6.2, Seurat v3.1.2) ^67,68^. The pipeline encompassed quality control, cell filtering, spectral clustering, cell type annotation, and visualization. Cells expressing 200 to 2500 genes that were detected in at least three cells and with mitochondrial gene content below 5% in droplets were considered viable. Cells of good quality from the five experimental groups were integrated for analysis using Seurat FindIntegrationAnchors and IntegratedData functions^67,68^. Principal component analysis (PCA) over the identified 2000 highly variable genes <u>were</u> applied for data dimension reduction (dimensions = 35) prior to cell clustering. Cell clustering was performed on integrated data with a shared nearest-neighbor (KNN) graph-based method using the FindNeighbors function included in Seurat, followed by the Louvain algorithm for modularity optimization (resolution = 2.5) using FindClusters function. After the cell clusters were determined, their marker genes were identified with the FindMarkers function. For cluster annotation, the top marker genes based on the adjusted p-value were manually curated to match canonical cell types and their marker genes, as identified in the literature research.

In scRNA-seq data P150-C1, raw single-cell RNA sequencing data was preprocessed using CellRanger v6.1.2 (10x Genomics), including aligning reads to mouse reference genome, and generating cellxgene expression count matrices. Subsequently , an in-house scRNA-seq pipeline including quality control, cell filtering, spectral clustering, cell type annotation, and visualization was implemented based on the Seurat R package (R v4.1, Seurat v4.1.1) ^67,68^. Cells expressing 200 to 6000 genes that were detected in at least three cells, with mitochondrial gene content below 20%, and with ribosomal gene content below 20% in droplets were considered viable. Cells of good quality from the two experimental groups were integrated for analysis using Harmony (v0.1.0)^69^. Principal component analysis (PCA) over the identified 2000 highly variable genes were applied for data dimension reduction (dimensions = 30). Subsequently, cells are assigned labels using the symphony package (v0.1.0) ^70^, with the P83 data serving as the reference. Symphony identifies the most similar cells in the project of reference and assigns the appropriate labels and UMAP position to the query sample.

### Cell-cell communication analysis

We used CellChat (version 1.4.0) along with its provided database of ligands and receptor pairs for mice to identify patterns of cell-to-cell communication. To assess the likelihood of communication, we followed the methodologies outlined in the original research paper by Jin et al. ^71^. We applied these methods at both the ligand-receptor pair level and the pathway level. To ensure reliable and significant findings, communication between cell types observed in fewer than 10 cells with adjusted p-values greater than 0.1 was excluded. The plots were generated using the built-in functions within CellChat.

### Pseudo time analysis using Monocle3

We used the Monocle3 v1.0.0 package as a Docker image from Biocontainers to perform pseudotime analysis to infer cell differentiation trajectories^72,73^. Gene expression counts, cell and gene metadata from scRNAseq samples were loaded as Monocle3 new_cell_data_set object. Data was pre-processed using 30 principal components to compute the sequence of gene expression changes and removed batch effects by using the sample id as an alignment group. Next, we generated a UMAP low-dimensional representation of the data using the reduce_dimensions function and fitted a graph describing the developmental process, ensuring all clustered partitions were joined. Pseudotime was generated by specifying a root node at the Alpha cells and visualized as a UMAP to observe the developmental progress through the projected trajectory. To ensure consistency with original UMAP low-dimensional representation, initial computed coordinates were transferred to the Monocle3 object prior to visualization.

### Gene-concept network analysis

To investigate genes involved in diabetes-related pathways, gene-concept network diagrams were generated using cnetplot function provided by the clusterProfiler R package (version 4.6.2)^74^. Specifically, genes such as *FOXO1, IGF2, Igf2r, CTCF, epn2, Pdgfra, Vegfa, Kdr, Hifa1, Flt4, Ulk1* are highly expressed in the pathways of interest, such as insulin-like growth factor receptor signaling pathway, gene expression by genomic imprinting, IGF signaling, and insulin receptor signaling pathway, as well as the top significantly enriched Gene Ontology (GO) terms, are also shown in this network.

### Bulk RNA-seq analysis

We used trimmomatic (Version 0.39) ^75^ to trim the low-quality next generation sequencing (NGS) reads (-threads 20 ILLUMINACLIP:TruSeq3-PE.fa:2:30:10 LEADING:3 TRAILING:3 SLIDINGWINDOW:4:15 MINLEN:36). Subsequently, only the high-quality trimmed reads were aligned to the mouse reference genome using STAR (Version 2.7.2b)^76^. The read counts were calculated by featureCounts software (Version 2.0.3)^77^. Differentially expressed genes (DEGs) were identified by using the DESeq2 R package (Version 1.26.0) (adjusted p-value <0.05)^78^. KEGG pathways and gene ontology (GO) enrichment tests were performed by the clusterProfiler R package (Version 3.14.3). A pathway or GO term was treated as significantly enriched if an adjusted p-values (with Benjamini-Hochberg correction) was smaller than 0.05. All bar plots and dot plots illustrating significant pathway or GO terms were created using the enrichplot R package (Version 1.6.1). Volcano plot of genes assigned colors based on fold change differences (>1.5- fold) and FDR-p values (p < 0.05). Statistical analysis was performed using the two-sided Wilcoxon test implemented by Seurat.

### Mouse primary endothelial cells and HUVECs

The remaining cells isolated from the pancreas used for scRNA-seq as described above were seeded and incubated in the complete endothelial cell growth medium: 40% F-12K medium (Corning), 40% DMEM medium (Corning), 20% fetal bovine Plasma (Neuromics), 1% home-made bovine brain food, 0.2% Heparin (Sigma-Aldrich, 50mg/mL), 1% penicillin-streptomycin (ThermoFisher Scientific), and 0.1% ciprofloxacin (Corning, 12.5 mg/mL) in 35-mm dish plates coated with 0.2% gelatin for about three days. Cells were washed daily with PBS to remove floating dead cells and debris, and the live cells attached to the plate were continuously incubated in fresh EC medium. When the cell confluency was around 70%, cells were harvested and purified with anti-mouse CD31 microbeads (Miltenyi Biotec, 130-097-418) according to standard manufacturer’s instructions. Briefly, cells were dissociated with trypsin-EDTA, then suspended in endothelial cell culture medium and centrifuged at 1,200 rpm for 5 minutes. The cell pellets of around 10^7^ cells were resuspended in 100 μL 1X binding buffer (PBS, pH 7.2, 0.5% BSA and 2 mM EDTA), then 10 μL anti-mouse CD31 microbeads were added into the cell solutions. The cell mixtures were incubated at 4^0^C in the dark. After 15 min incubation, the cells were washed with the binding buffer. The CD31^+^ endothelial cells were isolated by using an MS column and separator, and were seeded into pre-coated 0.2% gelatin cell culture plates immediately for the following experiments.

For dermal endothelial cells isolated from mouse skin, the back of the mice was shaved after sacrifice. The dorsal skins were isolated and washed with 70% ethanol and following washing with PBS. The skins were cut into smaller pieces on ice and incubated in around 10 mL digestion enzyme solution: 5 mg/mL Dispase II (Sigma-Aldrich, neutral protease, grade II), 2 mg/mL Collagenase Type I (ThermoFisher Scientific, 17-100-017), 0.01 g/mL BSA (Sigma-Aldrich, Fraction V) in DMEM (ThermoFisher Scientific, 11995073), incubated at 37^0^C for 1.5 h. The digested skins were then filtered through a 70-μm cell strainer. The cells were washed with pre-warmed DMEM two more times, and then incubated in the complete endothelial cell culture medium as described above. The subsequent purification steps using anti-mouse CD31 microbeads are the same as for endothelial cells isolated from pancreas. The process for isolating endothelial cells from mouse lung was almost identical to isolating endothelial cells from pancreas as described above. For mouse primary endothelial cells, passages 1 to 2 were used.

HUVEC cells from Lonza were cultured on 0.2% gelatin-coated plates in complete endothelial cell growth medium as described above, and cells at passages 5 to 10.

### Bulk RNA-seq library preparation

Primary endothelial cells isolated from pancreases of WT:STZ/HFD, EC-iDKO:STZ/HFD, EC-iFOXO1-KO:STZ/HFD, and EC-iULK1-KO:STZ/HFD mice were immediately used for RNA extraction. Total RNA extraction was performed using the RNeasy Minikit (Qiagen) following the manufacturer’s instructions. Values of RNA integrity number (RIN) were measured with an Agilent Bioanalyzer. The 2100 Bioanalyzer (Agilent) was used for quality control of all RNAseq samples in the Biopolymers facility at Harvard Medical School. RNA library was prepared using the TruSeq mRNA-seq mRNA-Seq Library Prep and sequenced using the HiSEQ Next-generation Sequencing System with a read length 75 at the core of AZENTA LIFE SCIENCES.

The following RNAseq raw data analysis read mapping and alignment, differential gene expression, gene-set enrichment analysis (GSEA), heatmaps were performed on the pipelines from the Bioinformatics Group at Boston Children’s Hospital and Harvard Medical School.

### Mass spectrometry analysis

Isolated and purified endothelial cells (EC) from WT:STZ/HFD, EC-iDKO:STZ/HFD, EC-iFOXO1-KO:STZ/HFD, and EC-iULK1-KO:STZ/HFD mice were incubated in the EC-conditioned medium (40% F-12K medium, 40% DMEM medium, 0.2% Heparin, 1% penicillin streptomycin, and 0.1% ciprofloxacin) for o/n. About 30 ml of cell culture supernatants were collected from three 10-cm dishes in which the EC cells were at approximately 80% confluency. The supernatants were further centrifugated at 1200 rpm for 10 min at 4^0^C or filtered through a 0.22-μm filter to remove the cell debris, and then concentrated into 300 μl using the Amicon® Ultra 15 mL centrifugal devices with Ultracel® 3KDa cutoff filters (Millipore). Experiments were repeated three times and biological samples were submitted to Taplin Mass Spectrometry Facility at Harvard Medical School for protein sequence analysis by LC-MS/MS. In brief, excised gel bands were cut into approximately 1 mm^3^ pieces and then subjected to a modified in-gel trypsin digestion procedure^79^. The digested peptides were extracted and reconstituted in 5 - 10 µl of HPLC solvent A (2.5% acetonitrile, 0.1% formic acid). Samples were run in a nano-scale reverse-phase HPLC capillary column^80^. A gradient was formed and peptides were eluted with increasing concentrations of solvent B (97.5% acetonitrile, 0.1% formic acid). Eluted peptides were subjected to electrospray ionization and then entered into an LTQ Orbitrap Velos Pro ion-trap mass spectrometer (ThermoFisher Scientific, Waltham, MA) to produce a tandem mass spectrum of specific fragment ions for each peptide. Peptide sequences (and hence protein identity) were determined by matching protein databases with the acquired fragmentation pattern by the software program, Sequest (ThermoFisher Scientific, Waltham, MA)^81^. All databases include a reversed version of all the sequences and the data was filtered to between a one and two percent peptide false discovery rate.

### Autophagosome formation analysis by Electron Macroscopy

The isolated and purified endothelial cells were seeded on coverslips in dish plates at 80% confluency. Cells were fixed in 1:1 diluted EC medium and routine fixative (5% glutaraldehyde, 2.5% paraformaldehyde, 0.06% picric acid, 0.2 M sodium cacodylate buffer , pH 7.4). After 1 h fixation at room temperature, cells were submitted to electron microscopy facility at Harvard Medical School for autophagosome formation analysis. Briefly, cells were washed three times in 0.1M cacodylate buffer, then postfixed in 1% Osmium tetroxide (OsO4)/1.5% Potassiumferrocyanide (KFeCN6) for 30 min. After washing and dehydration in grades of alcohol (5 min each: 50%, 70%, 95%; twice 100%), cells were embedded in plastic by inverting Epon/Araldite-filled gelatin capsules onto coverslips and polymerizing at 60°C for 24 h. The coverslips were removed by immersing the block in LN2 after polymerization. Approximately 80 nm thick sections were cut using a Reichert Ultracut-S microtome, placed on copper grids, stained with lead citrate, and examined using a JEOL 1200EX transmission electron microscope. Images were recorded with an AMT 2k CCD camera.

The parameter and quantitation of autophagosome has been described previously^82^. Samples were blinded and three individuals did measurements randomly from different cells to minimize the bias measurements. The cytoplasm areas were manually traced by freehand tool in image J. The number of autophagosomes or/and phagolysosomes in the cytoplasmic region was manually counted from each cell. To ensure accurate and reproducible values, measurements were repeated on a minimum of 18 cells for each sample. Increasing the number of samples (n) by expanding the number of cells quantified can decrease the variability, especially if significant variability is observed between individuals performing the analysis. The scale bar in each image represents 500 nm. This factor is equal to 500 nm/353 pixels. The ratio of the number of autophagosomes or/and phagolysosomes in the cytoplasmic area of cell:

Ratio = (Number/cytoplasmic pixel^2^)*(factor)^2^.

### Pancreas CD31^+^ and IGF2R^+^ cells analysis by Flow cytometry

The total cells isolated from the pancreas of WT:SZT/HFD, EC-iFOXO1-KO:STZ/HFD, EC:iDKO:STZ/HFD, and EC-iULK1-KO:STZ/HFD mice for scRNA-seq, as described above, were resuspended in 100 μL FACS buffer (1X PBS, 2% FBS, 2mM EDTA). The cells were incubated with the primary antibodies: CD31 (1:100, eBioscience^TM^, 550274 from rat) and IGF2R (1:100, Cell Signaling Technology, 14364 from rabbit) respectively on ice for 30 minutes. Afterward, they were washed with 100 μL FACS buffer, spun down, and resuspended in 100μL FACS buffer containing fluorescent secondary antibodies: donkey anti-rat 488 (1:100, Invitrogen, A-21208) for CD31^+^ cells, donkey anti-rabbit 594 (1:100, Invitrogen, A-21207) for IGF2R^+^ cells. Following 30 minute incubation, the cells were washed with FACS buffer, fixed with 4% paraformaldehyde (PFA), and resuspended in FACS buffer for analysis. Single-color and no-color controls were prepared using ECs, which were treated the same as experimental groups. The expression of cell markers was analyzed using FlowJo version 10 software. Gating strategies were performed as described in the Figures.

### β -Cell culture

β-TC-6 (CRL-11506™), derived from (C57BL/6J × DBA/2J) F2 RIP1Tag2 mouse strain, a model of T2D was purchased from the American Type Culture Collection (ATCC, VA, USA), and maintained in the ATCC-formulated Dulbecco’s Modified Eagle’s medium (ATCC, 30-2002), supplemented with 15% FBS (Neuromics, FBS007), in 5% CO2 at 37 °C.

### Apoptosis assay

Three independent apoptosis assays were conducted: 1). β-cells (β-TC-6), cultured in 24-well plates to 70% confluence, were treated with 0 pg/mL, 1.5 pg/mL, 3 pg/mL, and 6 pg/mL IGF2 (R&D Systems). After 24 hours of incubation, the cells were harvested for apoptosis analysis; 2). β-cells (β-TC-6) were initially cultured overnight in 24-well plates to achieve 40% confluence. Subsequently, they were co-cultured with ECs isolated from WT:STZ/HFD, EC-iFOXO1-KO:STZ/HFD, EC-iDKO:STZ/HFD, EC-iULK1-KO:STZ/HFD mice (β-cells : ECs ratio approximately 1:1). After 24 hours of incubation, all cells were harvested for apoptosis analysis; 3). β-cells (β-TC-6) were initially cultured overnight in 24-well plates to attain 40% confluence. They were then co-cultured with ECs from the mentioned mouse groups (β-cells : ECs ratio approximately 1:1). After a 3-hour incubation, 1.5 pg/mL IGF2 (R&D Systems) was added. Following an additional 24-hour incubation, all cells were harvested for following apoptosis analysis.

For the apoptosis assay in 1), The β-TC-6 cells were incubated in 24-well plates with 40 % confluence for o/n, then added the same number of ECs isolated from pancreas of WT:STZ/HFD, EC-iFOXO1-KO:STZ/HFD, EC-iDKO:STZ/HFD, and EC-iULK1-KO:STZ/HFD mice (The ratio of β-cells : ECs was approximal at 1:1). After a 3 h incubation, 1.5 pg/mL of IGF2 (R&D Systems) was added into the cells. Following a 24 h incubation, all cells were harvested for the following apoptosis assay.

For the apoptosis assay in 2), cell apoptosis analysis was conducted by using the FITC Annexin V apoptosis detection Kit with 7-AAD kit (BioLegend, 640922) and flow cytometry (BD, the Accuri C6 Plus). Annexin V is a protein that binds to phosphatidylserine (PS) in a calcium-dependent manner. PS is translocated to the outer leaflet of cell surfaces during apoptosis and this is used as an indicator of cell apoptosis. Cell permeability dye of 7-ADD can detect the death cells. Hence, incorporating of 7-AAD with Annexin V staining via flow cytometry is a fast and reproducible way to detect different stages of cell apoptosis. To investigate IGF2-induced β-cells apoptosis and the influence exerted by ECs on apoptosis induced by IGF2, the cell mixture was stained with anti-CD31 (1:100, PECAM-1, eBioscience^TM^, 550274) to gate the CD31+ cells and separate the population of β-cells, which are CD31^-^ cells. By using this gating strategy, the β-cells can be properly gated on the cell populations shows a typical staining pattern on cells undergoing apoptosis using Annexin V and 7-AAD quadrant 1 (Q1) shows Annexin V and 7-AAD positive populations, which are at the terminal stage of cell death. Q2 shows Annexin V-positive and 7-AAD-positive populations, which are cells undergoing later apoptosis. Q3 represents Annexin V-positive and 7-AAD negative populations, which are early apoptosis cells.

For apoptosis analysis in 3), cell apoptosis was analyzed using the FITC Annexin V with 7-AAD Apoptosis Detection Kit (BioLegend, 640922) and flow cytometry (BD, Accuri C6 Plus). Annexin V binding to phosphatidylserine (PS) was indicative of apoptosis, while 7-AAD was used to detect death cells. To detect β-cells apoptosis induced by the combined effect of IGF2 with ECs, cells were first stained with anti-CD31 (primary, 1:100, eBioscienceTM, 550274 from rat), followed by staining with goat anti-rat 647 (secondary, 1:100, Invitrogen, A-21247), in combination with FITC Annexin V and 7-AAD. The gating strategy consisted of staining total cells to separate CD31^+^ cells (EC, stained and detected at wavelength 647) from CD31^-^ cells, which are β-cells. The quadrant analysis included Q1 (Annexin V and 7-AAD positive, representing the terminal stage of cell death), Q2 (Annexin V-positive and 7-AAD-positive, indicating cells undergoing later apoptosis), and Q3 (Annexin V-positive and 7-AAD negative, representing early apoptosis). The expression of cell markers was analyzed using FlowJo version 10 software. Gating strategies were performed as described in the Fig. 3d and Extended Data Fig. 6a, b, d.

### Immunofluorescence confocal microscope

The ECs or pancreatic tissues were fixed in ice-cold PBS containing 4% formaldehyde and permeabilized in PBS containing 0.2% Triton X-100, then blocked with blocking buffer (3% BSA, 3% donkey serum, 0.3% triton-100 in PBS) at room temperature for 1h. The sections were incubated with primary antibodies overnight at 4°C. The primary antibodies: rabbit anti-VEGFR2 (1:100, R&D Systems, AF357), anti-CD31 (1:100, PECAM-1, eBioscience^TM^, 550274), anti-VE-cadherin (1:100, R&D Systems, AF1002), anti-IGF2R (human, 1:100, R&D Systems, AF-2447-SP; mouse, 1:100, Cell Signaling Technology, #14364), anti-IGF2 (human, 1:50, R&D Systems, AF-292-SP; mouse, 1:100, ThermoFisher Scientific, PA5-78019), anti-CTCF (1:100, Santa Cruz Biotechnology, sc-271474), anti-LC3B (1:100, Santa Cruz Biotechnology, sc-398822), anti-ULK1 (1:100, Santa Cruz Biotechnology, sc-390904), anti-EEA1 (1:100, Santa Cruz Biotechnology, sc-6415), anti-LAMP1 (1:100, Santa Cruz Biotechnology, sc-18821), anti-epsin 1 (1:200, home-made). The primary antibodies were selected based on each staining markers of the cells or pancreas tissues.

The sections were washed three times by ice-cold wash buffer (0.3% triton-100 in PBS). The corresponding fluorescently labeled secondary antibodies were added and incubated at room temperature for 2 hours. The conjugated secondary antibodies were: donkey anti-rat 488 (1:200, Invitrogen, A-21208), donkey anti-mouse 488 (1:200, Invitrogen, A21202), donkey anti-rabbit 488 (1:200, Invitrogen, A-21206), donkey anti-mouse 594 (1:200, Invitrogen, A-21203), donkey anti-rabbit 594 (1:200, Invitrogen, A-21207), donkey anti-rat 594 (1:200, Invitrogen, A-21209), goat anti-rabbit 647 (1:200, Invitrogen, A-21244), goat-anti-rat 647 (1:200, Invitrogen, A-21247). The sections were mounted using mounting medium containing the nuclear dye 4’,6-Diamidino-2-phenylindole dihydrochloride (DAPI, 1:1000, Sigma-Aldrich, D9542-10MG). After thorough washing, the cells or tissues were mounted on coverslips. To verify antibody specificity and distinguish target staining from background, the isotypes containing only the secondary antibody were used as a control.

Immunofluorescence images were acquired using an Olympus 1X81 Disc Confocal Microscope equipped with an Olympus Plan Apo Chromat 60X (for Cells) or 20X (for Tissues) objectives and a Hamamatsu Orca-R2 Monochrome Digital Camera C1D600 for imaging. Tile scanning is used specifically for tissue imaging.

### qRT-PCR

Total RNA of ECs was isolated from different sources, including WT, WT:SZT/HFD, EC-iFOXO1-KO:STZ/HFD, EC:iDKO:STZ/HFD, and EC-iULK1-KO:STZ/HFD, or *db/db* mice using either commercial kits (Qiagen, Valencia, CA, USA) or TRIzol (Invitrogen) as per the manufacturer’s instructions, followed by DNase I treatment. Oligo(dT)20 primers (Invitrogen-18418-020) were used for reverse transcription of the extracted RNA into cDNA, and qRT-PCR was performed in triplicate using the StepOnePlus Real-time PCR system (Applied Biosystems, Foster City, CA, USA) and Platinum SYBR Green qPCR super mix (Invitrogen) and Assays-on-Demand Gene Expression probes (Applied Biosystems). The qRT-PCR primer sequences are listed as follows:

For mouse primers:

*FOXO1:* Forward, 5’-GCTTAGAGCAGAGTTCTCACATT-3’, 5’-CCAGAGTCTTTGTATCAGGCAAATAA-3’

*Igf2*: Forward, 5’ **-** GGTACCAATGGGGATCCCAG - 3’, 5’ - TCAGGGGACGATGACGTTTG - 3’;

*Igf2r*: Forward, 5’-GTTGGTGTAGGGCCAGTGTT-3’, Reverse, 5’-GCGGGGTACTTTGCTTTTGG-3;

*Ctcf*: Forward: 5’-GCCATTTTGTGTCCGAGCC -3’, Reverse: 5’-CCTTCCATCTCCCCTTCCTCT -3’;

*VE-cadherin:* Forward, 5’-TGCTCACGGACAAGATCAGC-3’, Reverse, 5’-GTGTTAGCATCGACCCCGAA-3’;

*Epn1*: Forward, 5’-AGATCAAGGTTCGAGAGGCAA-3’, reverse, 5’-GTGAGGTCGGCAATCTCTGA-3;

*Epn 2*: Forward, 5’-ACATCCCAGGTTTTAGGCCG-3’, Reverse, 5’-GGAGTTTGTGGTGGGGAGAG-3’;

*LC3B*: Forward, 5’-GCGAAGTGATTATTTCCCGC – 3’, Reverse, 5’-GCCTGAGACTACCTACCGGG -3’;

*ULK1:* Forward, 5’-CCCAGAGTACCCGTACCAGA-3’, Reverse, 5’-TGTAGGGTTTCCGTGTGCTC -3’;

*β-actin*: Forward, 5’-AGAGCTACGAGCTGCCTGAC-3’, Reverse, 5’-AGCACTGTGTTGGCGTACAG-3’.

For human primers:

*FOXO1:* Forward, 5’- GTGTCAGGCTGAGGGTTAGT-3’, Reverse, 5’-CTG CCA AGT CTG ACG AAA GG-3’;

*CTCF:* Forward, 5’ – AAGATGCCTGCCACTTACCC – 3, Reverse: 5’ – CCCTTCAGGCAAAGGTAGGG – 3’;

*KDr*: Forward, 5’-AAGGAGAAGCAGAGCCATGT – 3’, Reverse, 5’ – ACCGTACATGTCAGCGTTTG – 3’;

*LC3B*: Forward, 5’-GTCGGAGAAGACCTTCAAGC-3’, Reverse, 5’-AAGCTGCTTCTCACCCTTGT – 3’;

*BECN1*: Forward, 5’-CACTCAGCTCAACGTCACTG -3, Reverse: 5’ – CTGCCACTATCTTGCGGTTC -3’;

*Epn1*: Forward, 5′-GAGAGCAAGAGGGAGACTGG-3′, Reverse, 5’-GTGAAGACGTCAGCAAGGTC-3’;

*Epn2*: Forward, 5’ -CAGTCCCTCAACCCTTTCCT-3’, Reverse, 5’-CGAAGCTGGTTCAGTGTCAG-3’;

*Ulk1*: Forward, 5’-TCCAAACACCTCGGTCCTCT-3’, Reverse: 5’-AACTTGAGGAGATGGCGTGT-3’;

*β-actin*: Forward, 5′-AGCTGCTTCTGCGGCTCTAT-3′, Reverse, 5′-GTGGACAGTGAGGCCAGGAT-3′.

For the ULK1 knockdown in HUVECs, the siRNA transfection of HUVECs was performed using Lipofectamine RNAiMAX (Invitrogen) following the manufacturer’s protocol. The following target sequences were used: human FOXO1 (Bioneer, 1058762) or ULK1 (Forwars, 5’-G.C.A.U.G.G.A.C.U.U.C.G.A.U.G.A.G.U.U.U.U-3’, reverse, 5’-A.A.C.U.C.A.U.C.G.A.A.G.U.C.C.A.U.G.C.U.U-3’). The HUVECs were harvested 48–72 h after transfection.

### Co-immunoprecipitation analysis and Western blotting

The immunoprecipitation procedure involved lysing transfected HEK 293T cells with RIPA buffer, comprised of 1% Triton X-100, 0.1% SDS, 0.5% sodium deoxycholic acid, 5 mM tetrasodium pyrophosphate, 50 mM sodium fluoride, 5 mM EDTA, 150 mM NaCl, 25 mM Tris (pH 7.5), 5 mM Na3VO4, 5 mM N-ethylmaleimide, and a cocktail of protease inhibitors. Pre-clearing of the cell lysates with mouse IgG and protein G Sepharose beads (ThermoFisher Scientific) at 4°C for 2 h was carried out, followed by incubation with antibodies against Flag-epsin 1 (1:1000, Sigma-Aldrich, F3165) at 4°C for 4 h. Negative controls were performed using mouse IgG instead of specific antibodies. The precipitated proteins were eluted from the beads using 2% SDS in 50 mM Tris (pH 7.5) and were subsequently visualized by western blotting. The proteins were separated by SDS-PAGE (7.5% acrylamide), electroblotted onto a nitrocellulose membrane, and blocked with 5% milk (w/v). Anti-epsin 1 (1:10000, home-made), anti-epsin 2 (1:1000, Abcam, ab74942), anti-FOXO1 (1:1000, Cell Signaling Technology, #2880S), anti-IGF2R (1:1000, Cell Signaling Technology, #14364), anti-IGF2 (1:1000, ThermoFisher Scientific, PA5-78019), anti-CTCF (1:1000, Santa Cruz Biotechnology, sc-271474), anti-ULK1 (1:1000), anti-LC3B (1:100, Cell Signaling Technology, #83506S). The primary antibodies were incubated overnight at 4°C, followed by incubation with the corresponding horseradish peroxidase-conjugated secondary antibody for 1 h at room temperature. The immunoreactive proteins were detected using enhanced chemiluminescence with autoradiography. The plasmids for overexpression of FOXO1 pSELECT-HA-mFOXO1 and FLAG-FOXO1 (pCMV5) were purchased from AddGene. Adenoviral vectors expressing GFP were purchased from Vectors BioLabs (Philadelphia, PA). Transfection-ready VEGFR2-turbo-GFP (including mutants by site-directed mutagenesis) and their control DNA plasmids were obtained from OriGene (Rockville, MD). Quantification of Western blotting results was carried out using the National Institutes of Health (NIH) ImageJ software.

### Proliferation, tube formation, and migration

*In vitro* monolayer ECs isolated from the pancreas of WT:STZ/HFD and EC-iFOXO1-KO:STZ/HFD mice for proliferation, tube formation, and migration were conducted as previously described with modifications^7^. For proliferation analysis, primary ECs were cultured on gelatin-coated coverslips until they reached 70% confluence. Cells were serum starved o/n and then stimulated with/without 100 ng/mL VEGF-A (BioLegend, 583102) for 24 h. The DNA replication precursor analogue 5-Ethynyl-2’-deoxyuridine (EdU)-incorporated cells were detected the cell proliferation with the Click-iT EdU Alexa Fluor-555 Imaging Kit (Invitrogen, <u>C10338</u>) and combined with CD31 staining (1:100, BD Pharmingen, 553373), and imaging using a Zeiss LSM880 confocal scanning microscope. For migration assay, monolayer ECs with an approximate population of 1.2 X 10^5^ were cultured in 24-well plates, and initiated by scratching wound using a 200 μL pipet tip followed by two washes with EC media. After 8 h of serum starvation, the cells were changed with fresh media, and cultured with/without 100 ng/mL VEGF-A (BioLegend, 583102) for 12–24 hours. The migration of monolayer ECs was assessed by capturing three different images of wound areas from each well using a digital camera under a microscope (×4). The area of wound was then measured (mm^2^). For tube formation, primary pancreas ECs isolated from WT:STZ/HFD, EC-iFOXO1-KO:STZ/HFD mice. Approximately 2.0 X 10^5^ cells were cultured in 48-well plates, and coated with 300 μl Matrigel (BD). The cells were serum starved for o/n, and supplemented with/without 100 ng/mL VEGF-A (BioLegend, 583102). A branched EC network was visualized under a Leica phase-contrast microscope, and the area covered by the branched cells was measured (mm^2^). The quantification was analyzed by NIH Image J 1.60.

### Mouse corneal micropocket angiogenesis assay

Corneal micropocket assays were performed as previously described^7,83^. Briefly, mice were anesthetized (Avertin, 250 mg/kg), and sustained-release pellet containing 20 ng of VEGF-A (BioLegend, 583102) were implanted into the corneal stromal layer. Five days after implantation, corneas were harvested and stained with PE-conjugated anti-CD31 antibody (1:100, BD Pharmingen, 553373) to visualize limbal vessels. Quantification was performed using growth pixels calculated with Adobe Photoshop, as previously described^7^.

### Statistics

Data were plotted and all statistical analyzes were performed using GraphPad Prism. Sample sizes were not predetermined through statistical methods; rather, they were determined based on the researchers’ experience and pilot data. Figure legends to the experiments provide details on statistical tests and post hoc analyses. A significance threshold of p < 0.05 was applied, and exact p values were reported. In comparisons, for the three groups of STZ/HFD mice, Dunnett’s multiple comparison test was used, with the WT:STZ/HFD as the reference. For multiple comparisons between several groups, Tukey’s post hoc test was used. Two-group comparisons were performed using Student’s two-tailed *t*-test, whereas Fisher’s exact test was used for categorical data.

### Study approval

All experiments performed in animals were approved by the IACUC at Boston Children’s Hospital, Harvard Medical School, Boston, MA. Studies using human samples were performed according to approved human research guidelines at Boston Children’s Hospital, Harvard Medical School, Boston, MA.

## Acknowledgements

We thank Ross Tomaino from Taplin Biological Mass Spectrometry Facility at Harvard Medical School for mass spectrometry analysis, Maria Ericsson from Harvard Medical School Electron Micrscope core for electron microscope study, Xiaofeng Cai and Hiroko Kishikawa for technical support for confocal images, Yang Lee and Anna Voronova for assistance of HUVEC and mouse primary endothelial cell culture studies, Marsha A. Moses, and Bruce Zetter for comments and suggestions.

This study was partially supported by grants from National Institute of Health (NIH) R01 HL133216, R01 HL130845, R01 HL141853, and R01 HL162367… to H.C.; ; 17SDG33410868 and NIH 5 T32 HL 7917-21/22/23 to H.W.

## Author Contributions

H. W. and H.C. conceived the project. H.W., K.C., B.Z., Y.W.L., L.S., Q.M., A.J., D.O.H, X.G., Y.L., S.B., H. F., A.E.B., M. V. M., M. L., H.C. participated in experimental design, execution, and data analysis. S.W. B.W., B. S., Q.P., V. N., K.L. contributed to a part of animal studies. H.W. and Y.W. L. prepared samples for bulk RNA-seq. H.W., C.K., B.Z., D.W. performed the cDNA library preparation for scRNA-seq. D.L.S., Q.M., A. J., D.O.H., X.G., Y.L., S.R., K. C., H.W., H.C. analyses the data of scRNA-seq and bulk RNA-seq. H.W., B.Z. performed the GTT and ITT assay. H.W., S.B., Y.W.L., M.L. J.X. performed most of the immunohistochemistry and confocal imaging studies. H.W., V.N. performed Western blot analysis and GSIS assay. S.E.B. and S.S. helped with plasma cholesterol and triglycerides analysis. H.W., K.C. performed the flow cytometry and apoptosis assay. H.W., H.F., A.E.B., R.J.D’.A, .S.B. performed the *in vitro* and *in vivo* angiogenesis and wound healing assay. H.W., H.F., K.C. did the quantification and statistics analysis. H.W. and H.C. wrote the manuscript. H.W., D.B.C., S.E.R., Y.W.L., J.X., Q.M., L.S., K.C., M.D., S.B., H.C. reviewed and edited the manuscript.

## Data and material availability statement

All data generated or analyzed during this study are included in this published article (and its supplementary data and source data files). All biological materials are available upon request from the co-corresponding author.

---

**Extended Data Fig. 1.**
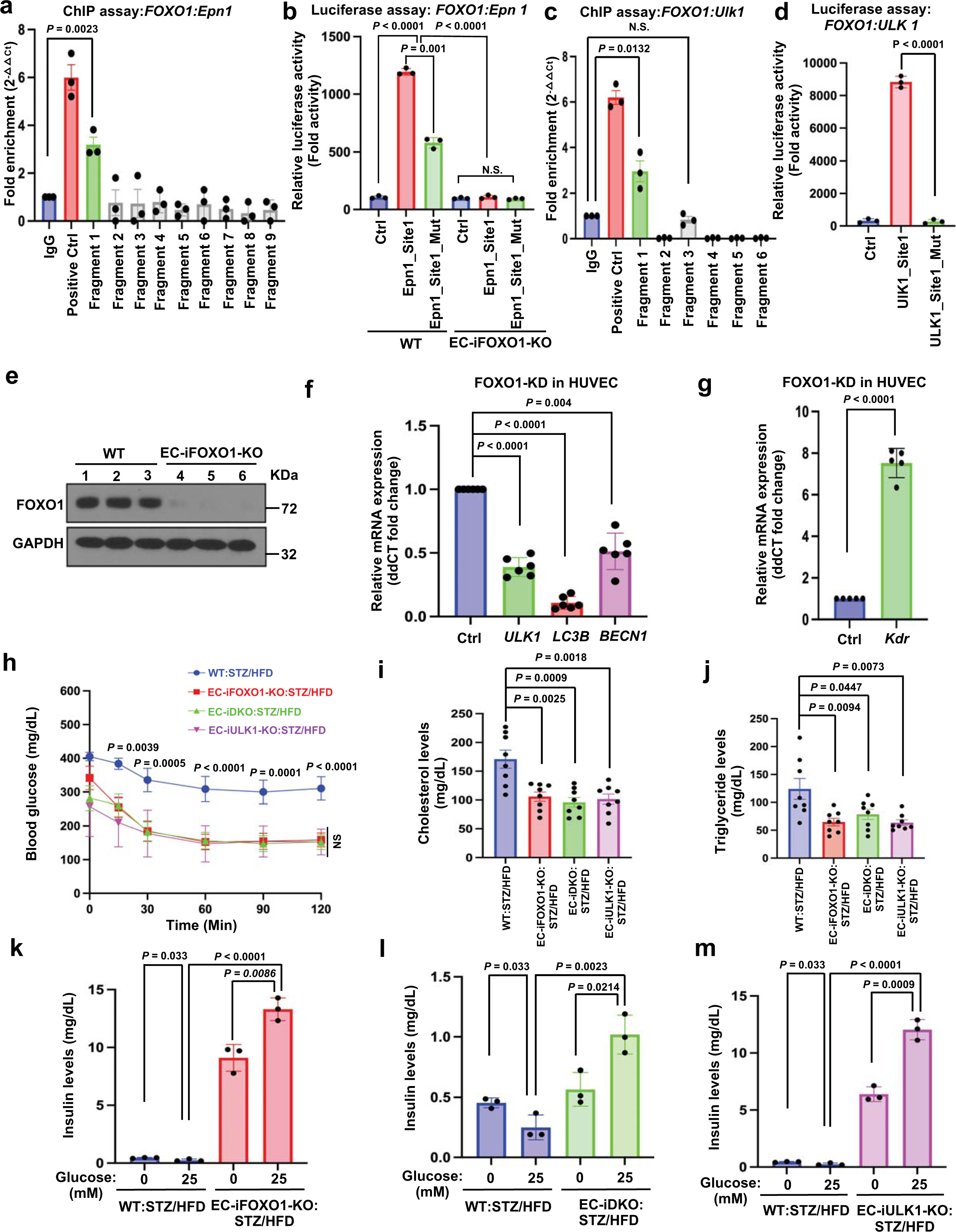
EC-specific knockout of FOXO1, epsins or ULK1 ameliorates β-cell dysfunction and improves diabetes. **a-d**, ChIP-qPCR assay (a, c) and luciferase analysis (b, c) indicate that FOXO1 can significantly bind to the promoter regions of the fragment 1, which includes the FOXO1 binding sequence element 5’-TGTTT-3’ in *Epn1* (a, b) and *ULK1* (c, d). n = 3. **e**, Western Blot shows FOXO1 knockdown in HUVECs (n = 3). **f, g**, qRT-PCR analysis of *ULK1*, *LC3B*, *BECN1*(**f**) and *Kdr* (g) expression in control and *FOXO1* knockdown HUVECs (n = 6 for *ULK1*, *LC3B*, *BECN1*; n = 5 for *Kdr*). **h**, ITT analysis showed that EC-specific knockout of FOXO1, epsins, and ULK1 in diabetic mice resulted in decreased fasting blood glucose levels after administrated 0.75 U/kg insulin compared with WT:STZ/HFD mice (n = 12). **i, j**, EC-specific knockout of FOXO1, epsins, and ULK1 in diabetic mice decreased cholesterol (**i**) and triglyceride (**j**) levels (n = 8). **k-m**, GSIS analysis shows insulin levels secreted by β-cells in islets from EC-iFOXO1-KO:STZ/HFD (**k**), EC-iDKO:STZ/HFD (**l**), EC-iULK-KO:STZ/HFD (**m**) mice were increased compared to WT:STZ/HFD mice after 25 mM glucose stimulation for 2 h. Islets from EC-iFOXO1-KO:STZ/HFD, EC-iDKO:STZ/HFD, and EC-iULK1-KO mice exhibited an increase in β-cell insulin secretion after exposure to 25 mM glucose for 2 hours compared to 0 mM glucose, while WT:STZ/HFD mice decreased in insulin secretion under the same conditions (n = 3). Statistical analyses in g, was 2-tailed Student’s *t*-test; for **a, b, c, d, k, l, m**, analyses were two-way ANOVA (P < 0.0001 overall) followed by Tukey’s post hoc test; in **h** was two-way ANOVA (P < 0.0001 overall) followed by Bonferroni’s multiple comparison test, comparisons for EC-iFOXO1-KO:STZ/HFD vs WT:STZ/HFD; **f, g, i, j**, analyses were one-way ANOVA (P < 0.0001 overall) with post-hoc Dunnett’s test, compared with WT:STZ/HFD control mice. Data in represent mean ± S.E.M. NS, not significant.

**Extended Data Fig. 2.**
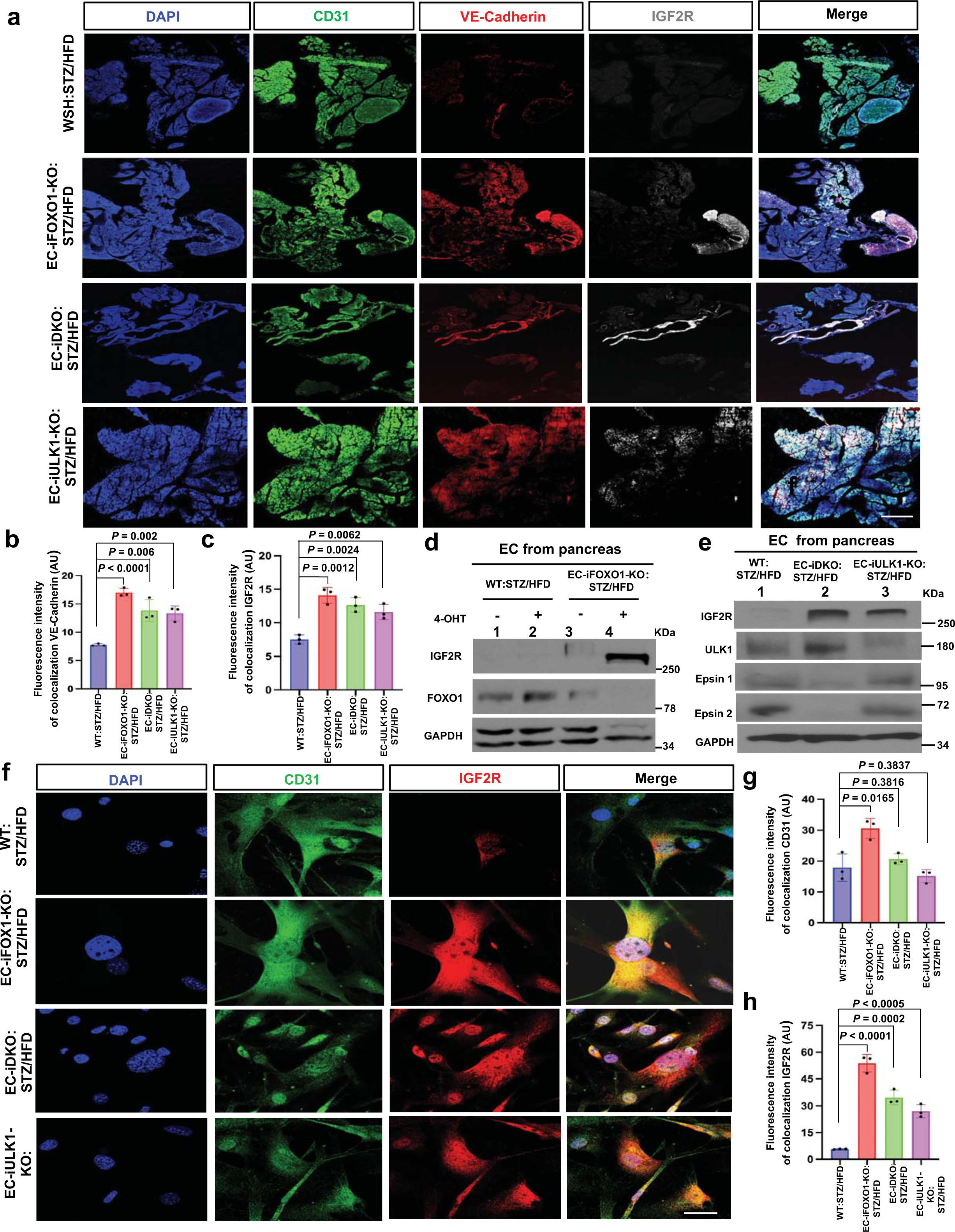
EC-specific knockout of FOXO1, epsins, or ULK1 increase IGF2R expression in diabetes. **a-c**, Representative confocal immunofluorescent tile scan images of the pancreas (**a**) and quantification of fluorescence intensities of VE-cadherin (**b**) and IGF2R (**c**) show that VE-cadherin and IGF2R levels were increased in EC-iFOXO1-KO:STZ/HFD, EC-iDKO:STZ/HFD, and EC-iULK1-KO:STZ/HFD mice compared with WT:STZ/HFD mice (n = 3). **d**, Western blot analysis shows IGF2R and FOXO1 levels in ECs isolated from the pancreas of WT:STZ/HFD and EC-iFOXO1-KO:STZ/HFD mice. ECs were treated with or without 4-hydroxytamoxifen (5 µg/ml) in EC medium. IGF2R expression was increased in EC-iFOXO1-KO:STZ/HFD mice compared with untreated ECs isolated from the pancreas of WT:STZ/HFD mice (N = 3). **e**, Western blot analysis of IGF2R expression levels in ECs isolated from the pancreases of EC-iDKO:STZ/HFD and EC-iULK1-KO:STZ/HFD mice showed an increase compared with WT:STZ/HFD mice (n = 3). **f-h**, Representative confocal immunofluorescence images of the ECs (**f**) and quantification of fluorescence intensities of CD31 (**g**) and IGF2R (**h**) show that CD31 and IGF2R levels were increased in EC-iFOXO1-KO:STZ/HFD, EC-iDKO:STZ/HFD, and EC-iULK1-KO:STZ/HFD mice compared with those in WT:STZ/HFD mice (n = 3). Statistical analysis in **b, c, g**, and **h** were one-way ANOVA (P < 0.0001 overall) with Tukey’s multiple comparisons test , compared with WT:STZ/HFD control mice. Data in represent mean ± S.E.M. Scale bars, 50 μm (**a**) and 20 μm (**f**).

**Extended Data Fig. 3.**
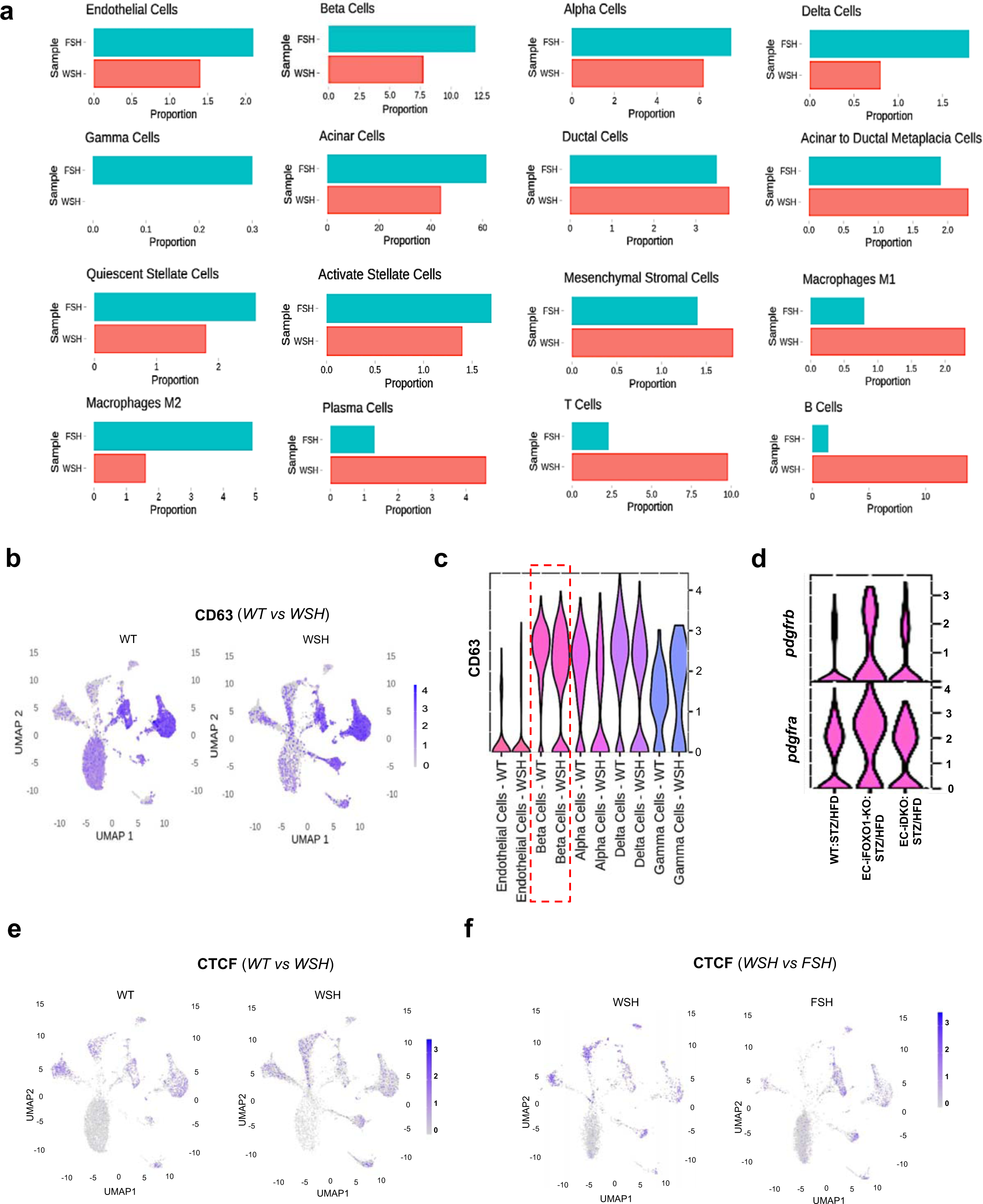
scRNA-seq shows a comparison of cell proportions between WT:STZ/HFD and EC-iFOXO1-KO:STZ/HFD, along with CD63 and CTCF expression in WT, WT:STZ/HFD, and EC-iFOXO1-KO:STZ/HFD. **a**, Comparison of pancreatic cell proportions in EC-iFOXO1-KO:STZ/HFD versus WT:STZ/HFD using scRNA-seq analysis. **b, c**, UMAP (**b**) and violin plot (**c**) show increased expression of *CD63* in WT compared with WT:STZ/HFD in β-cells. **d**, Violin plot showing increased expression of *Pdgfra* and *Pdgfrb* in EC-iFOXO1-KO:STZ/HFD and EC-iDKO:STZ/HFD mice compared with WT:STZ/HFD mice. **e, f**, UMAP shows increased *CTCF* expression in WT compared to WT:STZ/HFD (**e**) and increased *CTCF* expression in EC-iFOXO1:STZ/HFD compared to WSH (**f**).

**Extended Data Fig. 4.**
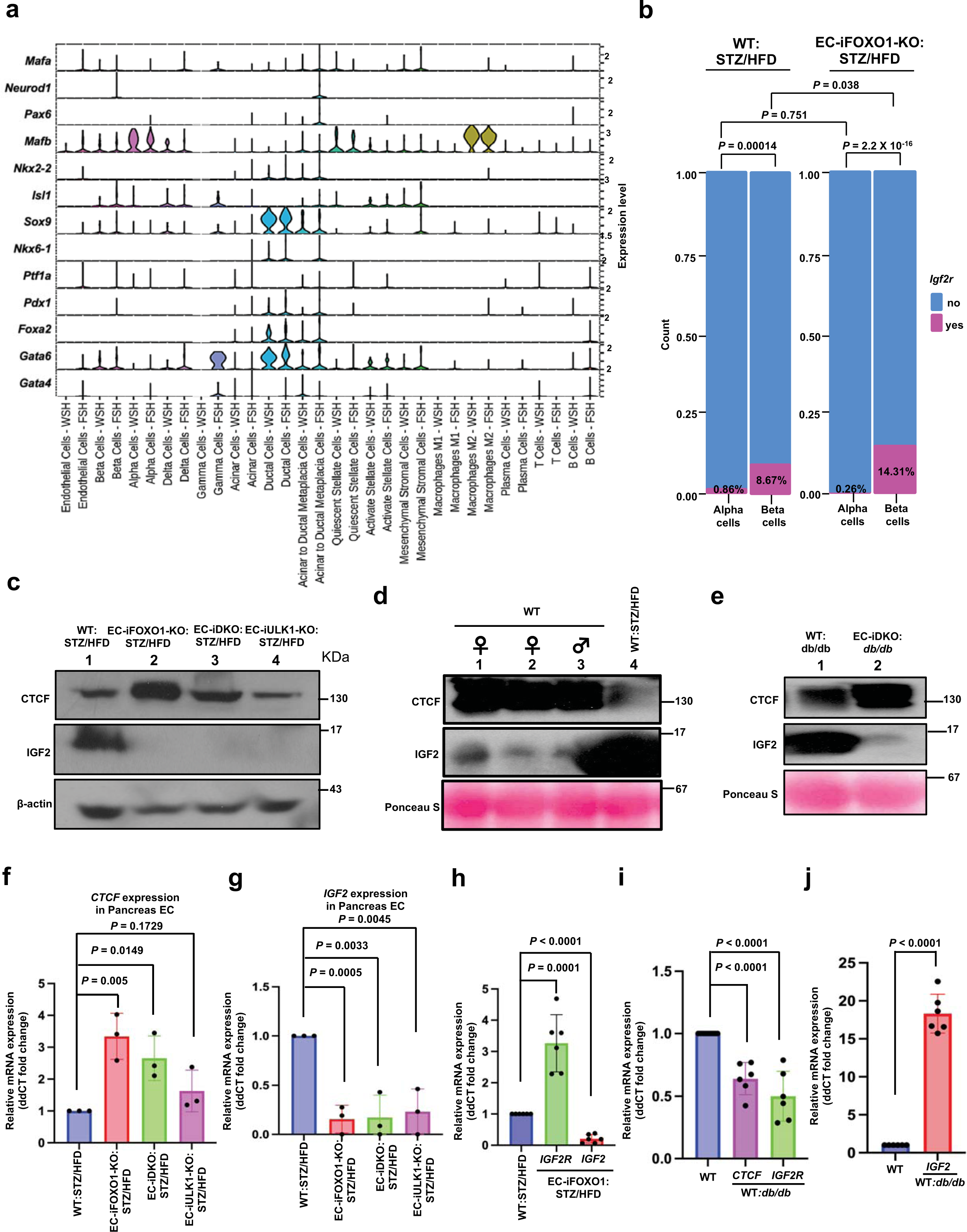
c-j supplement to Fig. 2d, e. FOXO1 regulates IGF2 expression through CTCF regulation and EC-FOXO1 depletion enhances IGF2R interaction activity and endocrine cell plasticity in the diabetic islet niche. **a**, Violin plots show increased expression of the transcription factors *Mafa*, *Mafb*, *Sox9*, and *Ptf1a* in ECs, β-cells, α-cells, and δ-cells, respectively, in EC-iFOXO1-KO:STZ/HFD compared to WT:STZ/HFD. **b**, The bar graph shows a higher proportion of cells expressing *Igr2r* in β-cells compared to α-cells. Moreover, approximately 14.31% of β-cells in EC-iFOXO1-KO:STZ/HFD expressed *Igr2r*, compared to 8.67% in β-cells of WT:STZ/HFD. Statistical analysis used one-sided proportion test. **c-e**, Western blot analysis showed increased CTCF levels and decreased IGF2 levels in skin endothelial cells isolated from EC-iFOXO1-KO:STZ/HFD, EC-iDKO:STZ/HFD, and EC-iULK1-KO:STZ/HFD compared with WT:STZ/HFD (**c**). CTCF levels were increased and IGF2 levels were decreased in the plasma of WT mice compared with WT:STZ/HFD mice (**d**). Furthermore, CTCF was increased while IGF2 was decreased in the plasma of EC-iDKO:*db/db* compared with WT:*db/db* mice (**e**) (n = 3). **f-j**, qRT-PCR analysis shows increased *CTCF* (**f**) and decreased *Igf2* (**g**) in EC-iFOXO1-KO:STZ/HFD, EC-iDKO:STZ/HFD, and EC-iULK1-KO:STZ/HFD compared with WT:STZ/HFD. *Igf2r* is increased and *Igf2* is decreased in EC-iFOXO1-KO:STZ/HFD compared to WT:STZ/HFD (**h**). Furthermore, *CTCF* and *Igf2r* were decreased (**i**), while IGF2 was increased (**j**) in WT:db/db compared to WT. For **f, g**, n = 3; for **h, i, j**, n = 6.

**Extended Data Fig. 5.**
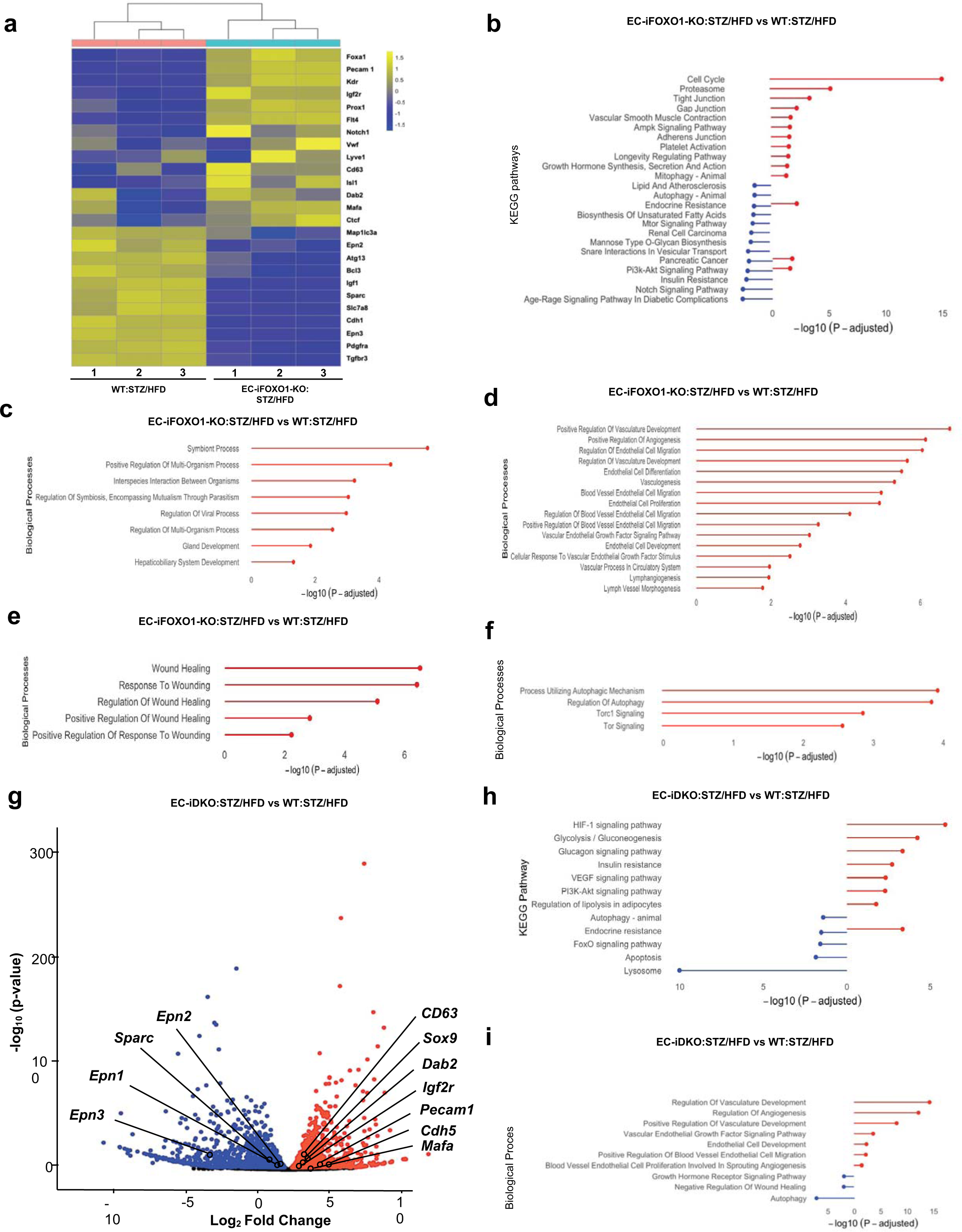
Comparison of Heatmaps and Gene Ontology (GO) Pathways from bulk RNA-seq Data: WT:STZ/HFD vs. EC-iFOXO1-KO:STZ/HFD and WT:STZ/HFD vs. EC-iDKO:STZ/HFD. **a-f**, Heatmap (**a**) of bulk RNA-seq shows genes, including *Igf2r, CTCF, Kdr, CD63, FOXA1*, were upregulated in ECs of EC-iFOXO1-KO:STZ/HFD compared with WT:STZ/HFD (n = 3). KEGG (b), Biological process (**c-f**) pathways between EC-iFOXO1-KO:STZ/HFD versus WT:STZ/HFD. **g-i**, Volcano plot showing the DEGs from ECs between EC-iDKO:STZ/HFD and WT:STZ/HFD. Genes, including *Igf2r, Prcam1, SOX9, CD63*, were upregulated in EC-iDKO:STZ/HFD compared with WT:STZ/HFD (**g**). The genes were color-coded based on fold change (>1.5-fold) and FDR-p values (p < 0.05). Statistical analysis employed the two-sided Wilcoxon test implemented via Seurat. KEGG (**h**) and biological process (**i**) pathways between EC-iDKO:STZ/HFD versus WT:STZ/HFD.

**Extended Data Fig. 6.**
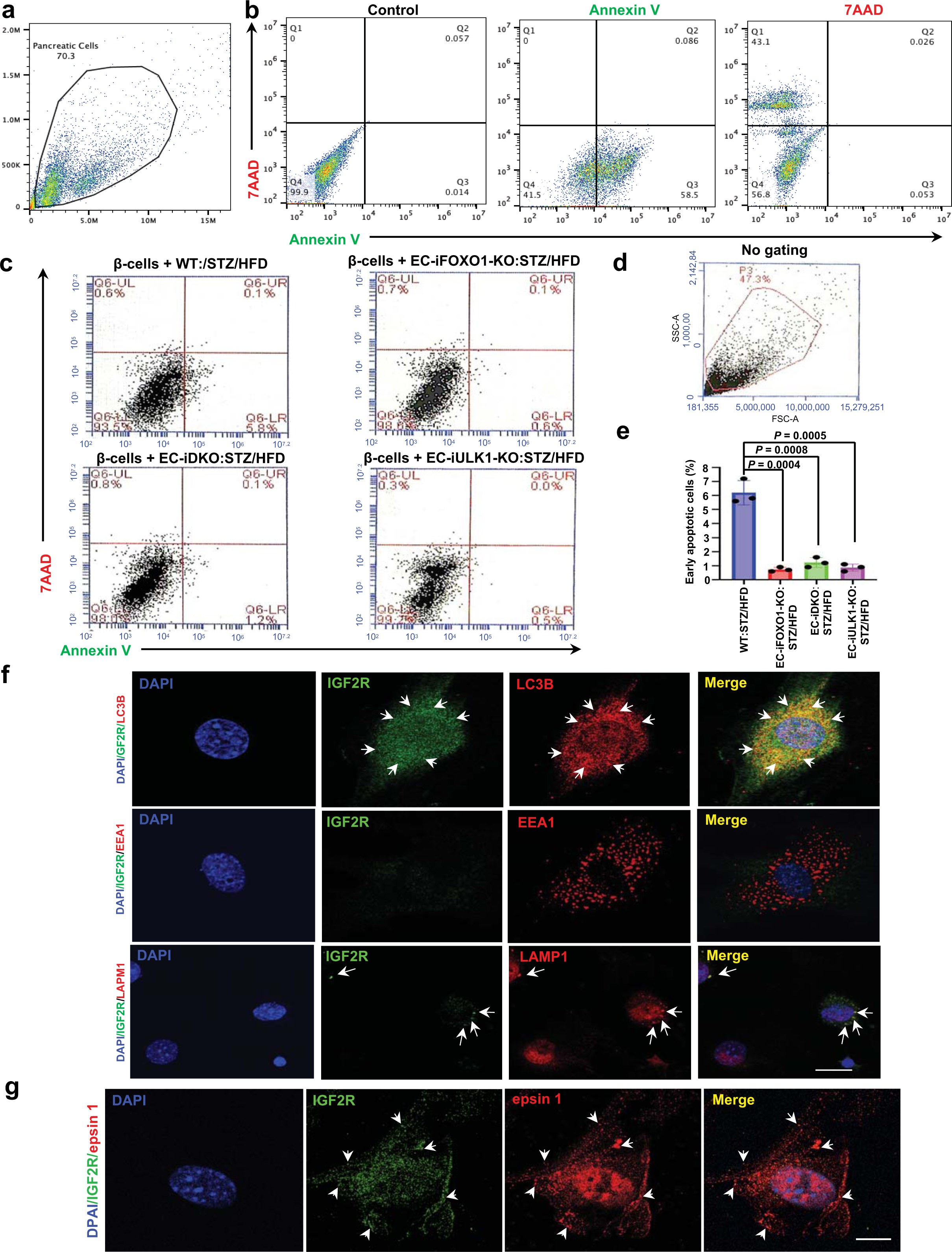
FOXO1, epsins, ULK1 regulate β-cell fate through the IGF2-IGF2R system in diabetes. IGF2 induces apoptosis of Beta-TC-6 cells apoptosis. **a, b**, Total pancreatic cells without gating (**a**). Gating strategy: quadrant 1 (Q1) shows both Annexin V- and 7-AAD-positive populations, which are at the terminal stage of cell death. Q2 shows Annexin V-positive and 7-AAD-positive populations, which are cells undergoing later apoptosis. Q3 represents Annexin V-positive and 7-AAD-negative populations, which are early apoptosis. **c-e**, Representative FACS plots of β-TC-6 cells co-cultured with ECs from pancreases of WT:STZ/HFD, EC-iFOXO1-KO:STZ/HFD, EC-iDKO:STZ/HFD, and EC-iULK1-KO:STZ/HFD(**c**), No gating cells (**d**), Quantification of percentages in Q6-LR (Annexin V-positive and 7-AAD-negative populations) of early apoptosis in c. (n = 3). **f, g**, Representative confocal immunofluorescent images of colonization of IGF2R with LC3B, EEA1, LAPM1 (**f**) and with epsin 1 (**g**) in the pancreas ECs isolated from WT:STZ/HFD mice (n = 3). Statistical significance in e was one-way ANOVA with Tukey’s multiple comparisons test, compared with WT:STZ/HFD control mice. Data in represent mean ± S.E.M. Scale bars: 20 μm.

**Extended Data Fig. 7.**
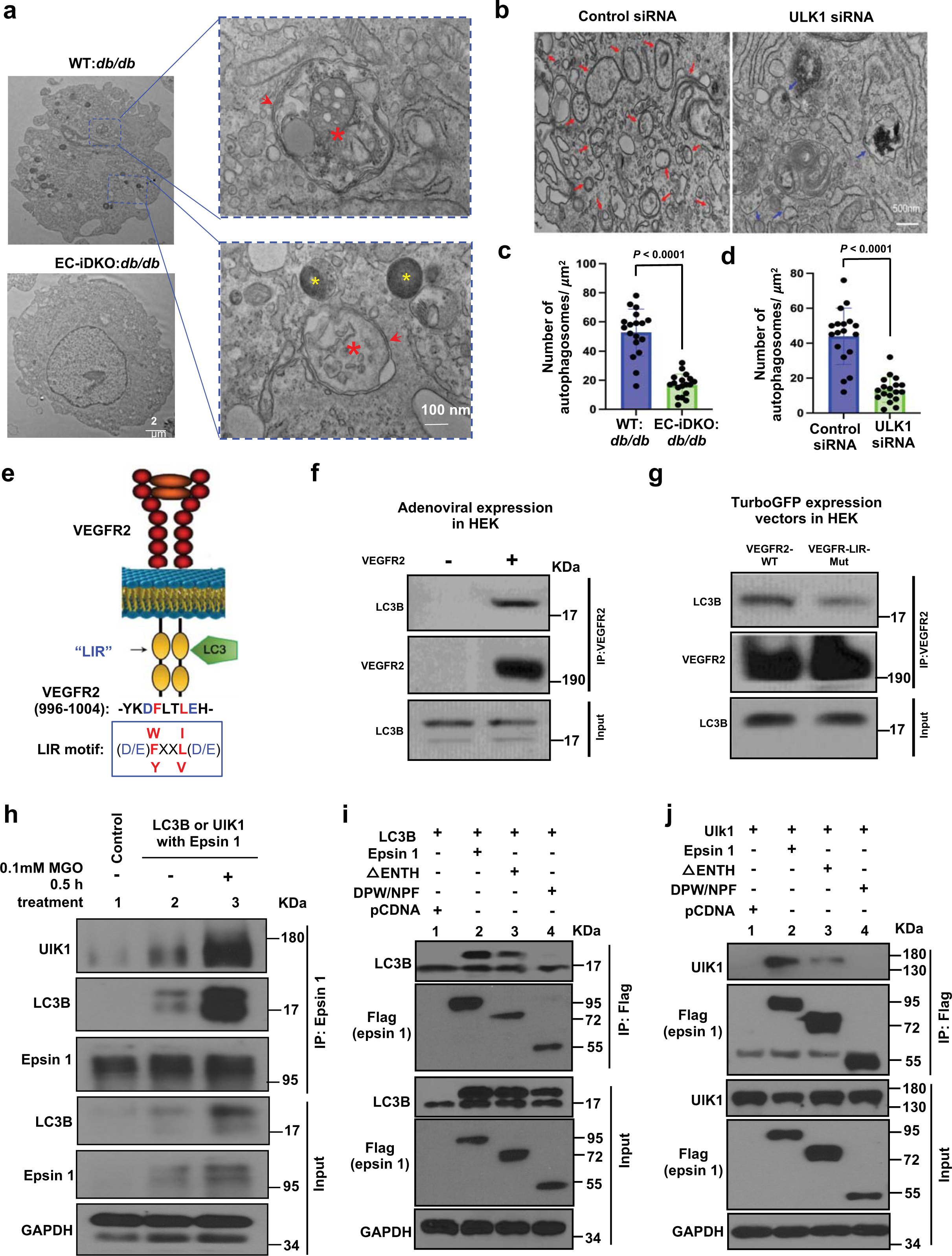
Epsins and ULK1 promote endothelial autophagosome formation and drive selective degradation of VEGFR2 in diabetes. **a-b**, Representative electron microscopy (EM) images showing electron-dense vacuoles of double-membrane autophagosome (red arrows) in the primary ECs of pancreases isolated from WT:*db/db* and EC-iDKO:*db/db* mice (**a**), ECs isolated from WT:db/db mice treated with control siRNA and ULK1 siRNA (**b**). (n = 18) **c**, Quantification of autophagosome in (**a**). **d**, Quantification of autophagosome in (**b**). **e**, Schematic showing LC3B binding motif of LIR “-YKDFLTLEH-” located in the kinase domain of VEGFR2. **f, g**, Co-immunoprecipitation demonstrates the interaction between VEGFR2 and LC3B (**f**), while blocking the interaction by mutating the LIR binding motif (**g**) in HEK cells. (n = 3). **h-j**, Co-immunoprecipitation of endogenous ULK1 and LC3B with epsin1 in mouse pancreas EC cells after 0.5 h stimulation with 0.1mM MGO stimulation (**h**). Full length and domain-deletion constructs of FLAG-epsin 1 within the pcDNA3 vector were co-transfected with ULK1 (**i**) or LC3B (**j**) into HEK 293T cells, followed by IP and western blot analysis using antibodies against FLAG (epsin 1) tags, as well as specific proteins (n = 3). Statistical significance in c, d was analyzed by a 2-tailed Student’s *t* test. Data in represent mean ± S.E.M. Scale bars 2 μm (a), 500 nm (b).

**Extended Data Fig. 8.**
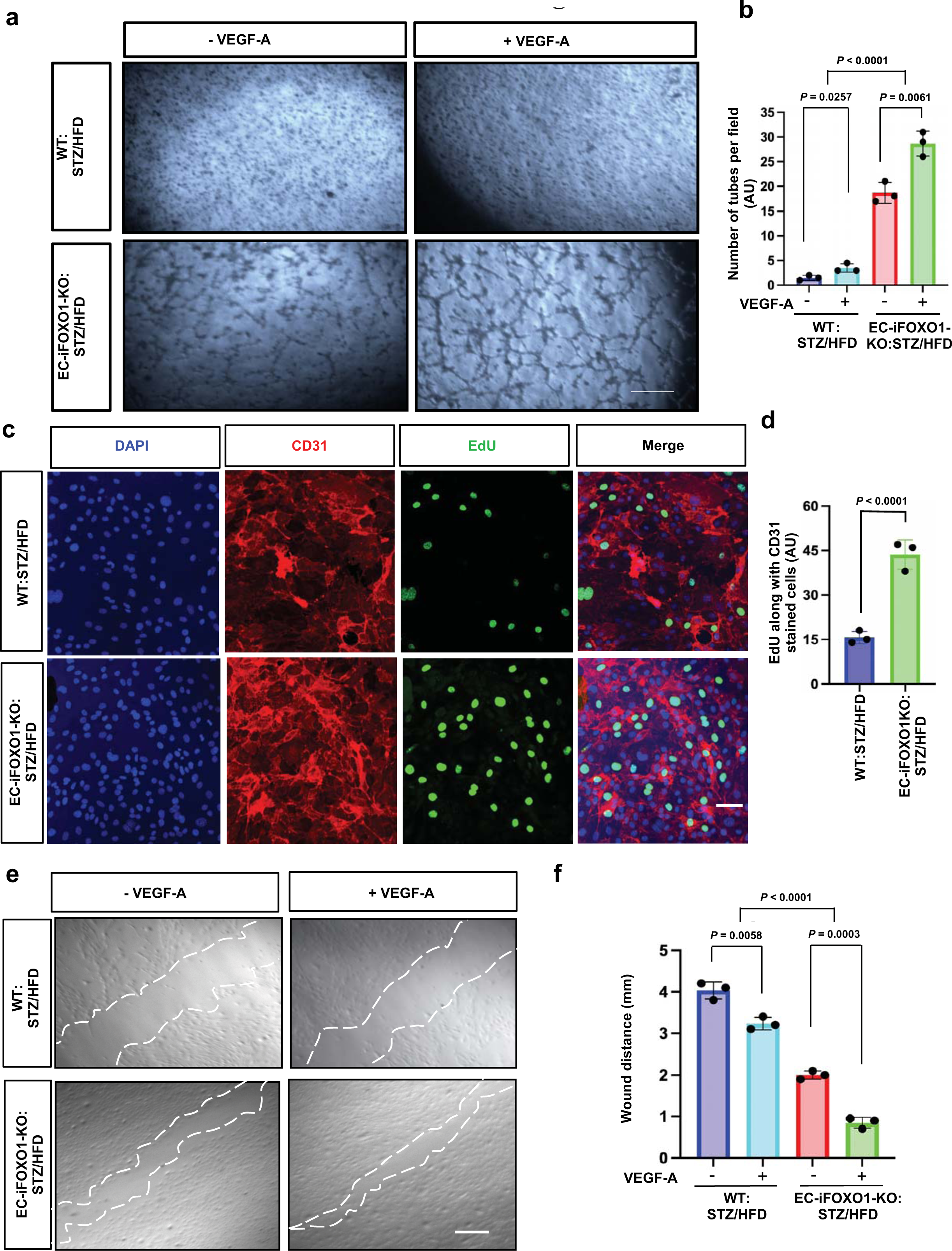
*In vitro,* EC-specific knockout of FOXO1 increases endothelial cells tube formation, proliferation and migration in diabetes. **a, b**, Representative images of pancreatic ECs isolated from WT:STZ/HFD and EC-iFOXO1-KO:STZ/HFD, at a concentration around 2 X 10^5^ cells/ml, were subjected to a tube formation assay by culturing on Matrigel in the absence or presence of 100 ng/mL VEGF-A for approximately 3 to 9 h (**a**). Quantification of tube formation is shown in (**b**). **c, d**, Representative immunofluorescence staining of CD31 (red) and EdU (green) in pancreas ECs isolated from WT:STZ/HFD and EC-iFOXO1-KO:STZ/HFD mice after treatment with 100 ng/mL VEGF-A for 24 h (c), and quantification of EdU along CD31 staining is shown in (**d**). **e, f**, Representative images of pancreatic ECs isolated from WT:STZ/HFD and EC-iFOXO1-KO:STZ/HFD LECs at 70% confluence were then subjected to a scratch assay in the absence or presence of 100 ng/mL VEGF-A for 16 h (**e**), and quantification of wound distance is shown in (**f**). The wound distance was image using a digital camera under a microscope (×4), the tube branched network was captured by using a Leica phase contrast microscope. For proliferation was imaged by a Zeiss LSM880 confocal microscope. The quantification was analyzed by NIH Image J 1.60. N = 3, all experiments were conducted at three independent biological experiments. Statistical analyses in **b, f**, were two-way ANOVA (P < 0.0001 overall) with Tukey’s multiple comparisons test. Significance in **d** was analyzed by a 2-tailed Student’s *t* test. Data in represent mean ± S.E.M. Scale bars: 50 μm (**a, e**), 20 μm (c).

**Extended Data Fig. 9.**
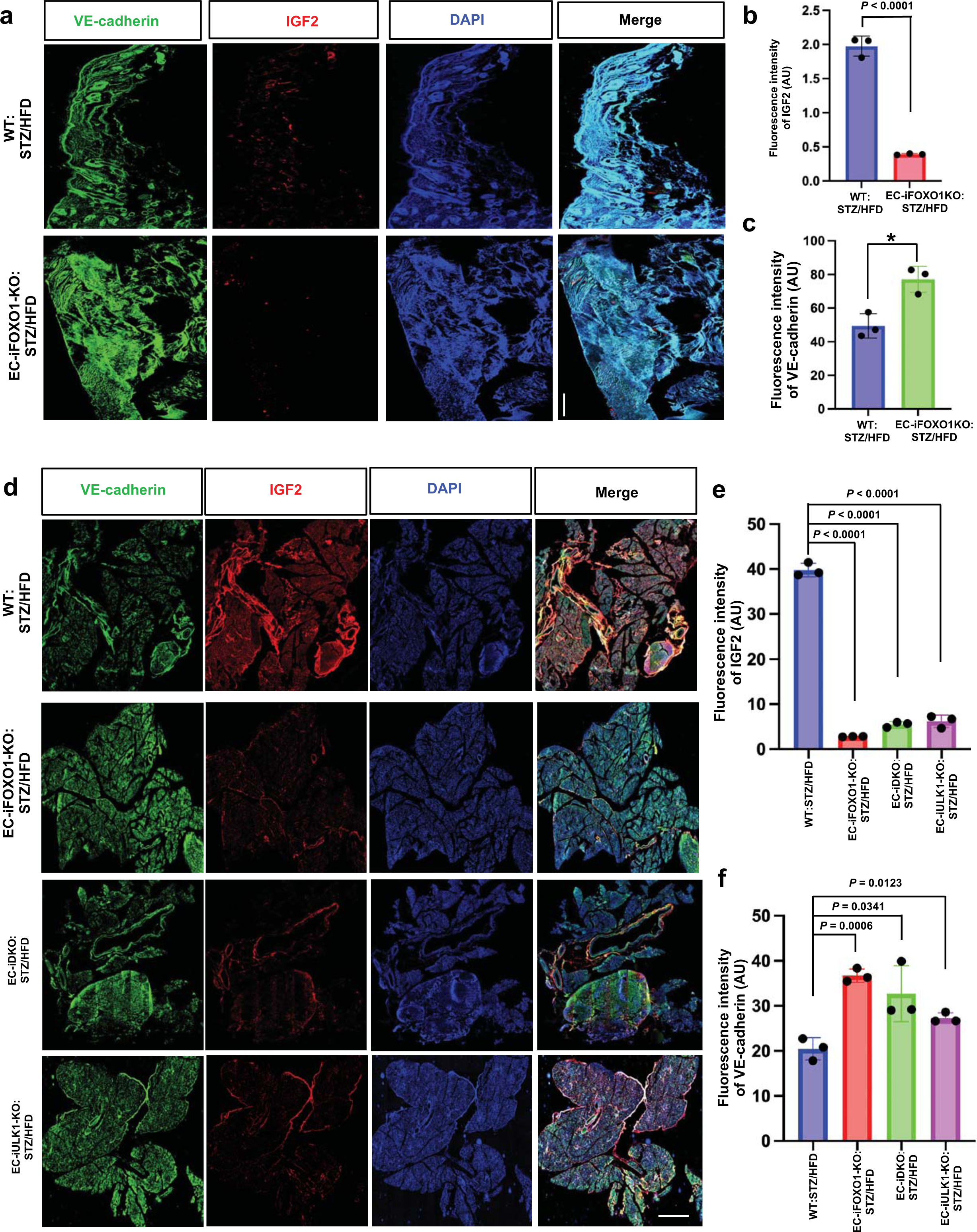
Immunofluorescence staining showed IGF2 expression in the pancreas and wounded skin in diabetic STZ/HFD mice. **a-c**, Representative confocal immunofluorescent tile scan images of the skin from WT:ST/HFD and EC-iFOXO1-KO;STZ/HFD mice were staining VE-cadherin (Green) IGF2 (red) (**specific knockout of FOXO1, epsins, and ULK1 ameliorates** ) and quantification of fluorescence intensities of IGF2 (**b**) and VE-cadherin (**c**). IGF2 levels were increased along VE-cadherin in WT:STZ/HFD compared with EC-iFOXO1-KO:STZ/HFD mice (n = 3). **d-f**, Representative confocal immunofluorescent tile scan images of the pancreas (**d**) and quantification of fluorescence intensities of IGF2 (**e**) and VE-cadherin (**f**) show that IGF2 levels were decreased in EC-iFOXO1-KO:STZ/HFD, EC-iDKO:STZ/HFD, and EC-iULK1-KO:STZ/HFD mice compared with WT:STZ/HFD mice (n = 3). Statistical analyses in **b**, **c** were analyzed by a 2-tailed Student’s *t* test. **e, f**, were one-way ANOVA (P < 0.0001 overall) with Tukey’s multiple comparisons test, compared with WT:STZ/HFD control mice. Data in represent mean ± S.E.M. Scale bars: 100 μm (**a**), 50 μm (**d**).

**Extended Data Fig. 10.**
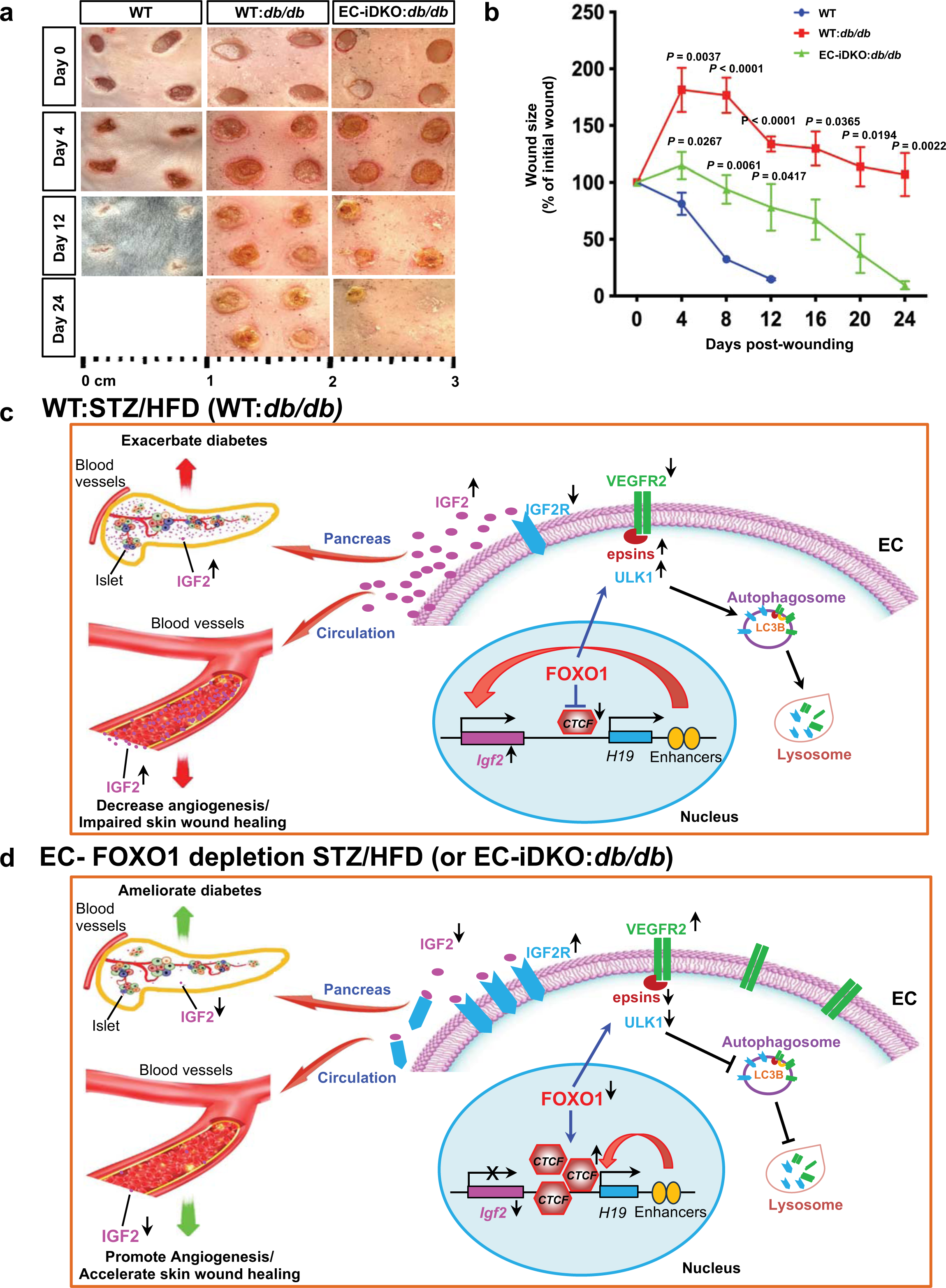
Wound healing of WT:*db/db* and EC-iDKO:*db/db* mice, and proposed mechanism by which EC-FOXO1 depletion reduces β-cell decline and improves wound healing in diabetes. **a, b**, Representative images of dermal wound healing at 0, 4, 12 days for WT, and at 0, 4, 12, 24 days for WT:*db/db* and EC-iDKO:*db/db* mice after dermal biopsy (n = 5). The quantification of wound area is shown in (**b**) and presented as wound healing curve. Quantifications were performed using NIH Image J. Significance in **b** was determined by two-way ANOVA followed by Bonferroni’s multiple comparison test, comparisons for EC-iDKO:*db/db* vs WT:*db/db* or EC-iDKO:*db/db* vs WT. Data are represented as means ± S.E.M. **c, d**, the proposed mechanisms by which EC-FOXO1 depletion reduces β-cell decline and improves wound healing in diabetes. FOXO1 inhibits CTCF and enhances IGF2 expression in diabetes. Meanwhile, FOXO1 upregulates epsins and ULK1, promotes IGF2R and VEGFR2 degradation through autophagosome formation, and ultimately inhibits angiogenesis. Hence, elevated IGF2 levels and reduced angiogenesis impair β-cell function, exacerbate diabetes, and impede wound healing in diabetes (**c**). In contrast, EC-FOXO1 deficiency increases CTCF levels and inhibits IGF2 expression in diabetes. EC-FOXO1 depletion prevents autophagosome formation and lysosomal degradation of IGF2R and VEGFR2, thereby promoting angiogenesis. Reduced IGF2 levels and enhanced angiogenesis are predicted to enhance β-cell function, improve diabetes, and accelerate skin wound healing (**d**).

